# Worm Perturb-Seq: massively parallel whole-animal RNAi and RNA-seq

**DOI:** 10.1101/2025.02.02.636107

**Authors:** Hefei Zhang, Xuhang Li, Dongyuan Song, Onur Yukselen, Shivani Nanda, Alper Kucukural, Jingyi Jessica Li, Manuel Garber, Albertha J.M. Walhout

**Affiliations:** Department of Systems Biology, University of Massachusetts Chan Medical School, Worcester, MA, USA; Bioinformatics Interdepartmental Ph.D. Program, University of California, Los Angeles, CA, USA; Via Scientific Inc. Cambridge, MA, USA; Department of Genomics and Computational Biology, University of Massachusetts Chan Medical School, Worcester, MA, USA; Department of Statistics and Data Science, Department of Biostatistics, Department of Computational Medicine, and Department of Human Genetics, University of California, Los Angeles, CA, USA

## Abstract

The transcriptome provides a highly informative molecular phenotype to connect genotype to phenotype and is most frequently measured by RNA-sequencing (RNA-seq). Therefore, an ultimate goal is to perturb every gene and measure changes in the transcriptome. However, this remains challenging, especially in intact organisms due to different experimental and computational challenges. Here, we present ‘Worm Perturb-Seq (WPS)’, which provides high-resolution RNA-seq profiles for hundreds of replicate perturbations at a time in a living animal. WPS introduces multiple experimental advances that combine strengths of bulk and single cell RNA-seq, and that further provides an analytical framework, EmpirDE, that leverages the unique power of the large WPS datasets. EmpirDE identifies differentially expressed genes (DEGs) by using gene-specific empirical null distributions, rather than control conditions alone, thereby systematically removing technical biases and improving statistical rigor. We applied WPS to 103 *Caenhorhabditis elegans* nuclear hormone receptors (NHRs) to delineate a Gene Regulatory Network (GRN) and found that this GRN presents a striking ‘pairwise modularity’ where pairs of NHRs regulate shared target genes. We envision that the experimental and analytical advances of WPS should be useful not only for *C. elegans*, but will be broadly applicable to other models, including human cells.

---

Since the dawn of functional genomics, the transcriptome has proven to be one of the most powerful molecular phenotypes to connect genotype to phenotype^1–3^. While early work in yeast provided insights into the transcriptional responses to gene deletions^4,5^, similar large-scale and systematic studies in multicellular organisms have been lacking. Moreover, statistical analyses of large-scale, high-throughput genomics data suffer from technical biases and high false discovery rates (FDRs)^6^, e.g., many false positives in the identification of differentially expressed genes (DEGs). More recently, a method commonly referred to as Perturb-seq has developed which uses pooled CRISPR-based gene perturbation screens with single cell RNA-seq. This method has proven powerful in in cell-based functional screens to annotate gene function, identify genetic interactions, and to infer disease-related pathways ^7–13^. However, this type of single-cell based approach is limited by the low sensitivity for gene detection, lack of biological replicate experiments and high cost, and challenges to perturb many genes *in vivo* ^3,14–16^.

Here, we present ‘Worm Perturb-Seq’ (WPS) in which individual genes are knocked down in the nematode *C. elegans* by feeding bacteria expressing double-stranded RNA, followed by RNA-seq using a strategy that adopts the high multiplexity of single-cell sequencing but uses bulk samples to produce high-resolution RNA-seq profiles. WPS is labor- and cost-efficient and enables replicate experiments. Using large-scale WPS data, we found that subtle but systematic fluctuations in gene expression caused pervasive false positive DEGs when analyzed by standard differential expression (DE) analysis methods that compare experiment to control conditions. To circumvent this issue, WPS includes a two-pronged data analysis framework, EmpirDE (‘<u>Empir</u>ical <u>D</u>ifferential <u>E</u>xpression’), that leverages the large WPS dataset to produce DEGs. EmpirDE systematically removes technical biases and uses gene-specific models with empirical null distributions to correct for anti-conservative *P* values (i.e., where the significance is overestimated) obtained by standard DE analysis. We demonstrate the rigorous control of FDR in WPS data and analysis by simulations and experimental benchmarking. We applied WPS to the knockdown of 103 nuclear hormone receptors (NHRs) and discovered that NHR pairs frequently share overlapping target genes, which cannot be explained by protein similarity, but is more related to NHR coexpression. WPS will enable examining different perturbations in addition to RNAi, including mutants, bacterial diets, and exposure to drugs or toxins. Importantly, EmpirDE will also broadly enable statistically rigorous analyses of large-scale transcriptomic data (i.e., > 100 conditions) in other systems.

## Results

In *C. elegans*, gene expression can be knocked down by feeding the animals bacteria expressing double-stranded RNA for a target gene of interest^17,18^. This whole-animal RNA interference (RNAi) is easy to perform for large sets of genes in parallel, and multiple RNAi libraries have been developed^19–23^. We developed WPS, which is composed of two major components: (1) an experimental approach to perform high-throughput whole-animal RNAi experiments and generate multiplexed RNA-seq libraries, and (2) a computational pipeline for quality control and rigorous statistical analysis of DEGs (**Fig. 1a**).

**Fig. 1:**
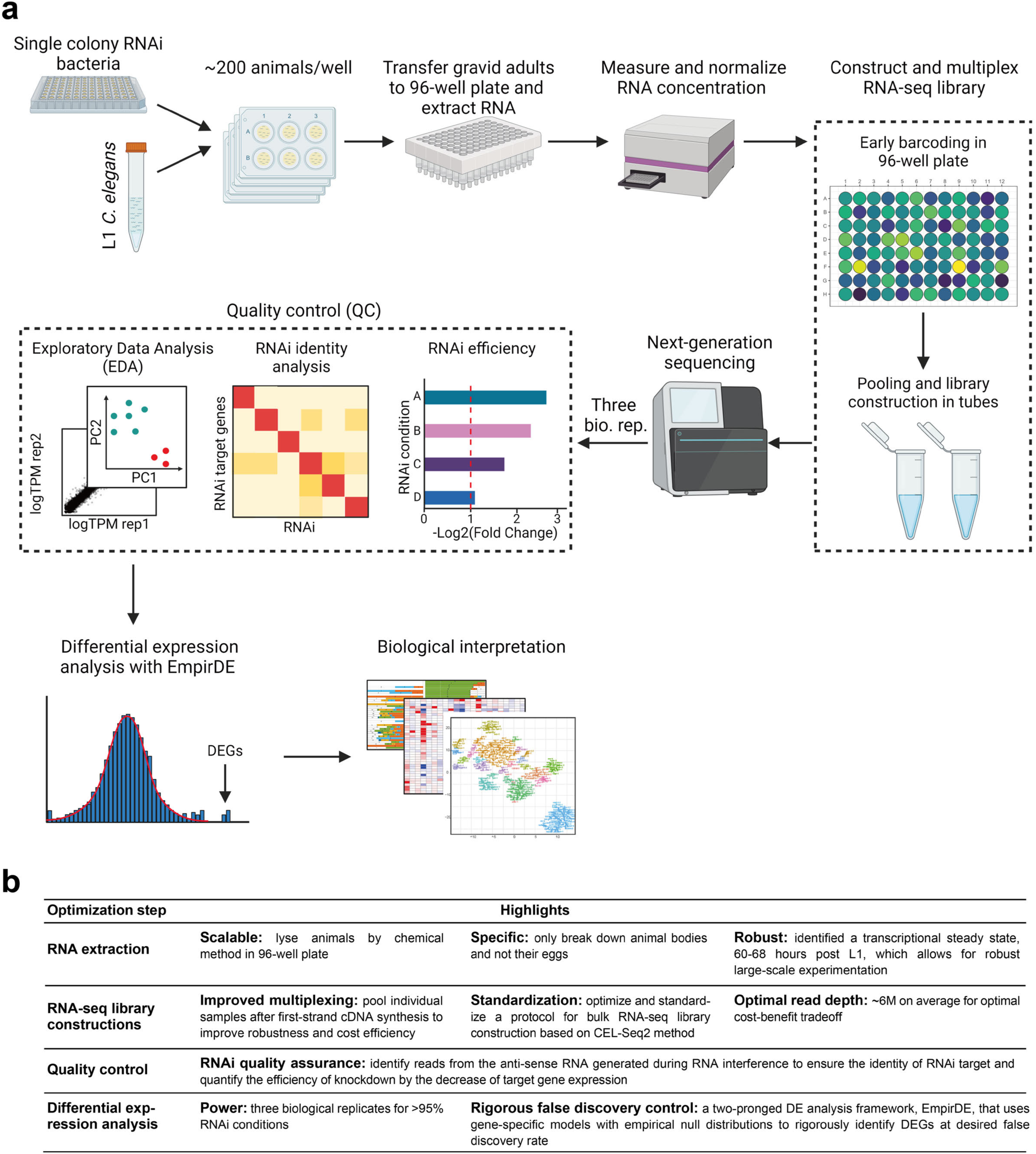
Worm Perturb-Seq (WPS) overview. **a**, Overview of the WPS pipeline. This figure was created with BioRender.com. **b**, WPS optimization highlights.

### An overview of Worm Perturb-Seq (WPS)

WPS consists of several steps, many of which were optimized to enable high-throughput, cost-effective experiments (**Fig. 1b**). Briefly, RNAi is started with animals at the first larval stage (L1) and, when grown to the desired stage, animals are harvested, and total RNA is extracted in a 96-well extraction plate. This streamlined workflow allows efficient triplicate experiments for hundreds of knockdowns (**Fig. 1a**). Multiplex RNA-seq libraries are constructed using an early barcoding step during reverse transcription in the 96-well plates, with each barcode linked to a single perturbation, followed by pooling of ∼50 samples and sequencing library construction. After sequencing, several quality control steps are performed (see below) and DEGs are identified with EmpirDE, which uses gene-specific models with empirical null distributions.

### An experimental WPS platform

Several experimental steps of WPS were developed and optimized, including growing animals, harvesting RNA, and generating, pooling, and sequencing of multiplexed libraries (**Fig. 1b**, **Supplementary Protocols** and **Supplementary Note 1**). Notably, WPS introduces a high-throughput worm lysis method for RNA extraction in 96-well plates, which does not lyse eggs, making it suitable to use WPS for gravid adults (**Extended Data Fig. 1a**). We optimized the 3’ end barcoding method CEL-Seq2^24^, which was originally developed for single-cell RNA-seq^25^, for multiplexing bulk RNA-seq libraries, with significant reduction of costly reagents (**Fig. 1b** and **Supplementary Protocol**).

The transcriptome of *C. elegans* changes greatly over the course of its lifetime; there are oscillatory expression profiles during development, and gene expression continues to change as the animals reproduce and age^26–28^. Therefore, if a knockdown has even a small effect on development, it can result in many differentially expressed genes that are secondary to the effect of the knockdown on development, rather than in response to the perturbed gene. We therefore opted to use a period in the animal’s lifetime in which the transcriptome does not change to minimize developmental effects of knockdowns. Animals develop from L1 to gravid adults in ∼58 hours at 20°C and we found that the gravid adult transcriptome was most stable between 60 and 68 hours post-L1-plating (**Fig. 2a, Extended Data Fig. 1b**). Therefore, we used a time of ∼63 hours post-plating, which is in between the first egg laid at 58 hours and the first egg hatched at 68 hours, allowing enough time for sample collection and processing, and providing a buffer for perturbations that elicit a mild developmental day (**Fig. 2a**).

**Fig. 2:**
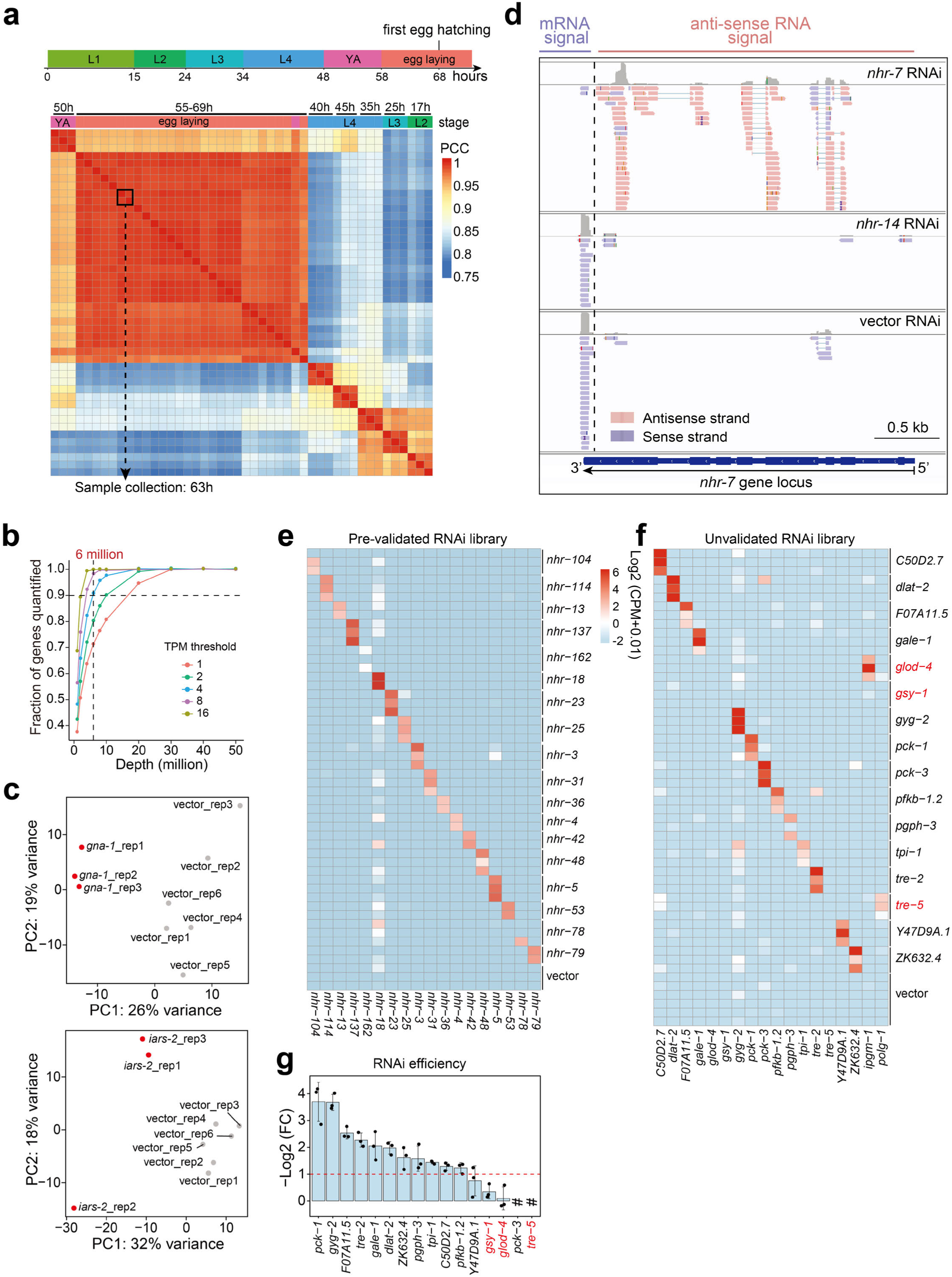
Development of WPS and data quality control. **a**, Comparison of the *C. elegans* transcriptome across developmental stages. The Pearson correlation coefficient (PCC) was calculated by the WPS profiles of animals fed vector control bacteria and collected at different time points post L1. **b**, Subsampling analysis of the reference WPS profile showing the fraction of genes quantified versus sequencing depth (see **Supplementary Methods**). **c**, Representative Principal Component Analysis (PCA) results for gene expression profiles of perturbations without (*gna-1* on top panel) and with (*iars-2* in bottom panel) a low-quality outlier replicate. Red and grey dots indicate RNAi and control samples, respectively. **d,** An example showing reads mapped to the reverse strand of the RNAi target gene (*nhr-7*) and a decrease in mRNA reads at the 3’ end. The reads mapped to *nhr-7* gene locus were visualized by Integrative Genomics Viewer (IGV ^70^). *nhr-7* RNAi was compared to vector control RNAi and another RNAi condition (*nhr-14*). **e, f**, Quantification of anti-sense RNA reads in a Sanger-sequenced (**e**) and an unvalidated (**f**) WPS sequencing library. Row names represent the intended RNAi gene (three replicates each) and column names represent the actual knocked down genes. Row names in red indicate wrong RNAi clones. Values are the log2(Count-Per-Million (CPM) + 0.01) of the reads mapped to the reverse strand of each gene in the columns. **g,** Log2(Fold Change (FC)) of the RNAi targeted gene expression for the WPS sequencing library shown in (**f**). Pound key (#) indicates not detected.

In other systems, it has been shown that most genes can be quantified with a relatively shallow read depth^29,30^. We performed down-sampling analysis of a reference dataset with a depth of 53 million reads (**Extended Data Fig. 1c and Supplementary Table 1**). We used an average of 6 million reads per sample, with which 90% of genes with >4 transcripts per million (TPM) and 80% of genes with > 2 TPM could be quantified (**Fig. 2b, Extended Data Fig. 1d**). For library multiplexing, we pooled ∼54 samples, which included 16 perturbations, each containing three biological replicates, together with six negative controls (empty vector RNAi) into one sequencing library. This design was intended to minimize batch effects by having all replicates of controls and perturbations in the same library. Here, we combined WPS datasets for 103 NHRs (discussed below) with those for ∼900 metabolic genes [REWIRING][WIRING] for benchmarking analysis.

### WPS quality control

Analysis of large-scale and high-throughput functional genomics experiments can be complicated by batch effects and low-quality samples^6,31,32^. We followed standard practices to ensure the quality of individual samples^33,34^, and to identify and remove outlier replicates (**Fig. 2c**, **Extended Data Fig. 1e**, **Supplementary Methods**). We next developed two RNAi quality control (QC) analyses to verify the gene that was knocked down. First, the reduction of targeted gene expression can be directly read out. For instance, for 85% genes that are expressed at a high level (TPM ≥ 30), we found a > 2-fold reduction in their mRNA levels when knocked down (**Extended Data Fig. 1f**). Second, due to abundant reverse strand reads that map to the gene body of knocked down genes, the identity of the perturbed gene can be directly identified from the WPS data (**Fig. 2d**, red reads). These reads are likely derived from unspecific reverse transcription of the anti-sense RNA generated during the RNA interference process^35^. This latter RNAi identity verification is particularly useful for genes that are expressed at low levels and was able to verify the identity of almost all NHR RNAi clones that were also confirmed by Sanger sequencing (**Fig. 2e**). We next performed WPS using RNAi clones that were not confirmed a priori [REWIRING] and found three incorrect clones (**Fig. 2f, g**, indicated in red). Importantly, we could identify the actual target by mapping anti-sense sequences to the *C. elegans* genome (**Fig. 2f**, non-diagonal signals for the red RNAi conditions). Two clones that returned a hit in the search were subsequently confirmed by Sanger sequencing (**Extended Data Fig. 1g**). The other incorrect clone had a partial insert that did not target a transcribed gene and was considered a non-targeting perturbation (NTP). In the metabolic-gene WPS screen [REWIRING] we found that ∼13% samples had a wrong RNAi identity, including 67 NTPs (**Supplementary Table 2**), showing the necessity of RNAi QC in large-scale screens. Taken together, WPS data can be directly used to validate that the gene that has been knocked down and clones that are incorrect can simply be removed from the dataset or analyzed with the corrected target information.

### Standard DE analysis results in high false discoveries

The 67 NTPs from the metabolic-gene WPS study [REWIRING] should have zero differentially expressed genes and can therefore be used to evaluate the actual FDR in WPS. We initially conducted DE analysis with DESeq2^36^, by comparing each RNAi perturbation to vector controls from the same sequencing library. Surprisingly, this approach resulted in dozens of DEGs in both NTP and four randomly spike-in vector control conditions (**Fig. 3a**, *P_adj_* < 0.01, fold change (FC) > 2, collectively referred to as NTPs hereafter). This suggests a high level of false discoveries despite stringent filtering by estimated FDR and FC thresholds.

**Fig. 3:**
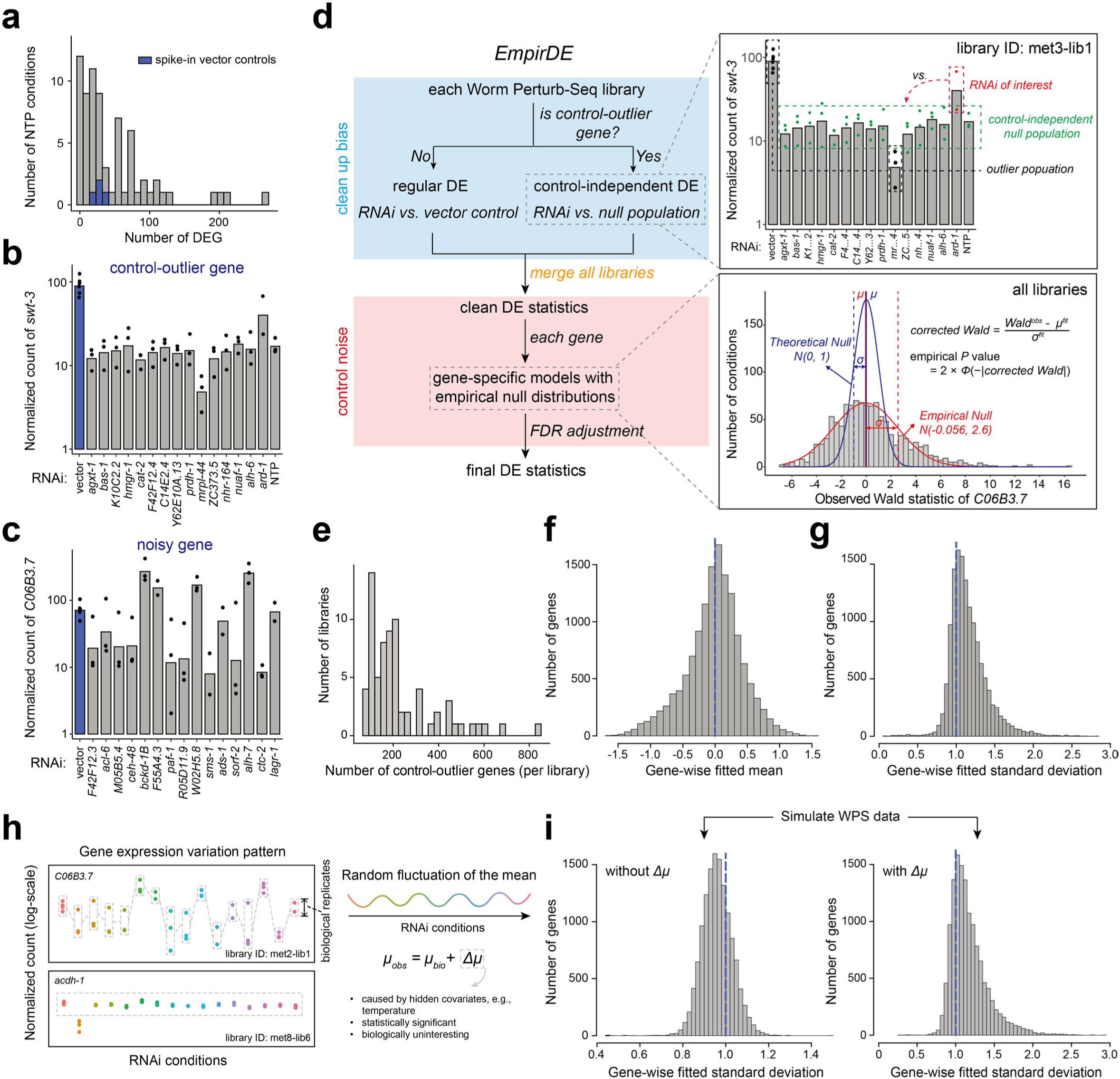
EmpirDE analysis framework reveals systematic anti-conservative *P* values caused by a deviation from the expected distribution of Wald test statistics. **a**, Distribution of the number of DEGs in non-targeting perturbations (NTPs) identified by DESeq2 analysis (FC > 2, adjusted *P* value (*P_adj_*) < 0.01). **b**, **c**, Examples of a control-outlier (**b**) and noisy (**c**) gene. Each bar plot shows the expression levels of the gene of interest in a WPS sequencing library that includes RNAi perturbations and vector control conditions. **d**, Schematic illustrating the EmpirDE framework. The zoom-in windows show two example genes shown in (**b**) and (**c**). **e**, The number of control-outlier genes per WPS sequencing library. **f**, **g**, Distribution of fitted means (**f**) and standard deviations (**g**) for all genes in the empirical null modeling. The blue dashed line shows the values for theoretical null. **h,** Examples showing the random fluctuation of the mean for two genes. **i,** Distribution of fitted standard deviation using simulated WPS data with (right) and without (left) adding a random fluctuation of the mean in the simulation.

False positive DEG calls are common in RNA-seq studies^37–40^ and are potentially more profound when combined with large-scale screens because of systematic variations^38^ ^6^. By comparing mRNA levels among vector control, NTP, and RNAi samples within the same sequencing library and across dozens of libraries, we discovered two confounding issues. The first issue involved sequencing libraries in which a gene consistently behaved differently in the vector control samples compared to the RNAi samples in the same sequencing library (**Fig. 3b**, ‘control-outlier gene’, **Extended Data Fig. 2a**). For instance, *swt-3* expression was lower in all RNAi samples when compared to the vector control in the same batch (**Fig. 3b**), resulting in *swt-3* being identified as a DEG in all these conditions, including in the NTP. The second issue involved genes with highly variable mRNA levels across the WPS dataset (**Fig. 3c**, see **Extended Data Fig. 2b** for the entire dataset, ‘noisy genes’) that were frequently called as DEGs in both RNAi perturbations and NTPs by DESeq2. This suggests that DESeq2 underestimates gene expression variances, leading to anti-conservative *P* values.

### EmpirDE: an empirical null-based, gene-centered method to rigorously analyze differential expression

A major challenge in transcriptomics is that successful DE analysis hinges on reasonable model assumptions and accurate parameter estimation. However, it is impossible to accurately estimate parameters, such as gene-specific variance, or to identify model misspecification, in small-sample-size experiments^41–44^. To systematically identify and correct for false positive DEGs, we developed EmpirDE, which leverages the power of having hundreds of conditions assessed in a uniform experimental setup, enabling the rigorous identification of true DEGs that are elicited by the specific knockdown. EmpirDE uses a two-pronged approach, first at the level of individual sequencing libraries and second at the level of an entire dataset (**Fig. 3d**). The first step performs DE analysis within each sequencing library (usually with ∼16 conditions) using DESeq2. However, instead of simply comparing an RNAi condition to the control, this step identifies control-outlier genes and treats these differently using a control-independent DE analysis procedure (**Supplementary Note 2**). Briefly, for each control-outlier gene, this step empirically identifies a control-independent null population based on the distribution of the gene’s expression levels in the sequencing library and compares the level for the gene in each RNAi condition to the newly defined null population. The second step of EmpirDE combines the DE results from all sequencing libraries (here ∼1,000 triplicate conditions) to correct the anti-conservative *P* values based on the gene-specific empirical null distributions of the DE test statistic (i.e., Wald statistic). Dozens to hundreds of conditions uniformly collected in WPS provided a unique opportunity to directly estimate the empirical null from the data in a gene-centered manner^45^. Assuming real effects are rare in large-scale experiments, the gene expression in most conditions can be viewed as ‘unchanged’, thus defining an empirical null population. By fitting the central peak of the distribution of the test statistic (i.e., the distribution of conditions that did not have a significant effect)^45^, this step estimates the empirical null and rescales the original test statistic (i.e., Wald statistic) accordingly to obtain a corrected Wald statistic on a gene-by-gene basis. This corrected statistic should follow a standard normal distribution and can be converted to a *P* value (referred to as empirical *P* value) to identify DEGs (**Fig. 3d**).

EmpirDE identified fewer than 200 control-outlier genes in most sequencing libraries (**Fig. 3e**), indicating a relatively low but significant number of confounded genes (1.4% of 14,000 detected genes). In the scenario of a well-fitted DE model, the empirical null distribution of the Wald test statistic in DESeq2 analysis should adhere to its theoretical null, a standard normal distribution^36^. Surprisingly, we observed a systematic difference between the theoretical and empirical null over the ∼14,000 detected genes in this dataset (**Fig. 3f, g**, **Fig. 3d** shows an example gene *C06B3.7*). While the mean of the empirical null distribution was symmetrically aligned around the mean of the theoretical Wald distribution (0) (**Fig. 3f**), the empirical null had systematically larger standard deviations than the theoretical expectation (1) (**Fig. 3g**). As a result, *P* values computed using the theoretical Wald distribution were anti-conservative, but this can be corrected by EmpirDE (**Fig. 3d**, empirical *P* value).

We next investigated the source of larger-than-expected standard deviations of empirical null. By inspecting mRNA levels across all conditions for each gene, we found a wide-spread random, yet consistent across samples, fluctuation of the mean (**Fig. 3h**, **Extended Data Fig. 2b**). This fluctuation is typically mild in its effect size, thereby distinguishing it from a specific change induced by RNAi (**Fig. 3h**, *acdh-1*). We hypothesized that the broad Wald statistic distribution is caused by such random fluctuations, and that the mean of observed gene expression (*μ_obs_*) is the sum of the actual biologically relevant expression change (*μ_bio_*) and a random experimental or condition specific fluctuation (*Δμ*) (**Fig. 3h**). The random fluctuation can be driven by systematic hidden covariates in the experiment, such as subtle difference in temperature associated with position in the culture plate, a common occurrence in large-scale experiments^6^. As such these technical fluctuations can be misinterpreted as biological signal from the RNAi treatment in WPS.

To test the hypothesis that a random fluctuation of the mean can introduce the observed test statistic inflation, we performed a simulation study using scDesign3^46^ to simulate the metabolic-gene WPS dataset [REWIRING]. To mirror the real data, we used the DEGs identified in the WPS analysis as the ground truth, and synthesized a new dataset in which the mean expression of each DEG was altered based on the DEG fold change while remaining consistent otherwise. To test the role of *Δμ*, we further introduced a random fluctuation to each gene based on the level of fluctuation of a gene in real data (**Extended Data Fig. 2c**). Consistent with our hypothesis, we found that the standard deviations of the empirical null distributions in simulated data matched the inflated test statistic of real data when a random *Δμ* was added, while being very close to the theoretical distribution when removing Δ*μ* (**Fig. 3i**, **Extended Data Fig. 2d-f**). Therefore, the observed anti-conservative *P* values can be explained with random fluctuations of mean mRNA levels.

### EmpirDE framework rigorously controls FDR

To benchmark the performance of EmpirDE, we first used simulation data to assess both FDR and power. As expected, both EmpirDE and conventional DESeq2 analysis had FDR correctly controlled on the simulated data without *Δμ* (**Extended Data Fig. 3a**, **b**). However, with *Δμ* added, only EmpirDE was able to rigorously control the observed false discovery proportion (FDP) at the expected FDR (**Fig. 4a**). Importantly, the power of EmpirDE is greater than that of DESeq2 at the same level of observed FDP (**Fig. 4b**), indicating a true increase in performance instead of nominal rescaling of *P* values. Such rigorous control of FDR depends not only on the optimized statistical modeling framework but also on the proper adjustment for multiple testing. WPS experiments involve both simultaneously testing the expression of thousands of genes in each perturbation (column-wise multiple testing) and hundreds of gene perturbations (row-wise multiple testing). Using the simulation data, we found that the worst-case adjusted *P* values in both column-wise and row-wise adjustments of a DE test aligned best with the expected FDR (**Fig. 4c**), while its loss of power was negligible (**Extended Data Fig. 3c**).

**Fig. 4:**
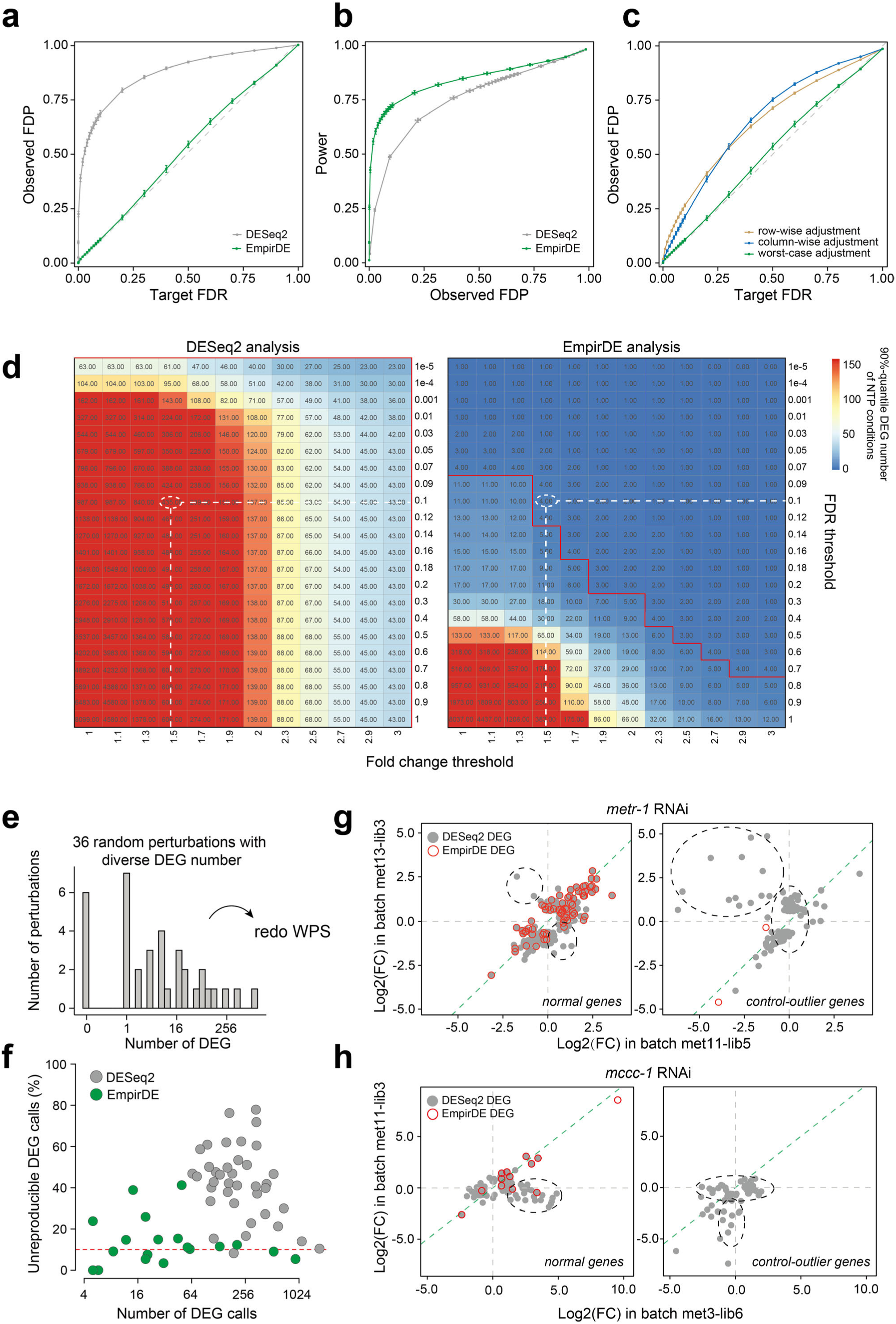
EmpirDE rigorously controls FDR. **a, b,** Benchmarking the performance of EmpirDE analysis framework. The observed False Discovery Proportion (FDP) is compared to target FDR (**a**) and power (**b**). The full metabolic-gene WPS dataset (3,691 samples) was simulated 10 times with random mean fluctuation (*Δμ*) to produce the error bars of each metric. FDP and power were measured based on a pooled set of 117,096 simulated DE changes in all conditions (**Supplementary Methods**). **c**, Benchmarking FDR control of different multiple testing adjustment strategies. **d,** Evaluating false discoveries using NTP experiments. The color (and numbers) in the heatmap shows 90%-quantile DEG numbers in the NTP experiments. The red lines show the threshold boundary for five false positive DEG calls. **e,** Number of DEGs (defined by FDR < 0.1 and FC > 1.5 using EmpirDE) for 36 perturbations that were repeated by a second WPS experiment. **f,** Fraction of unreproducible DEGs for EmpirDE versus DESeq2 analysis. Unreproducible DEGs were defined by genes that are called as DEG in one experiment (FDR < 0.1, FC > 1.5) but confidently non-DEG in the other (FC < 1.1 or show a different FC direction). The red dashed line shows the theoretical FDR (FDR = 0.1). **g**, **h**, Comparison of log2(FC) measured in two independent experiments for representative RNAi with either high (**g**) or moderate (**h**) number of DEGs. The green dashed line indicates the diagonal (y=x) and black dashed circle indicates genes that are not reproduced.

Next, we experimentally benchmarked the EmpirDE performance. We first used the NTP experiments mentioned above (**Fig. 3a**) to compare the number of false positive DEGs between DESeq2 and EmpirDE analysis with different thresholds for both FDR and fold change (**Fig. 4d**). We found that the 90% quantile of the number of DEGs detected in the NTPs was much lower with EmpirDE compared to DESeq2 (**Fig. 4d**). Specifically, at a FC of 1.5 and FDR < 0.1 we detected 4 and 435 false positive DEGs in the EmpirDE and DESeq2 analysis, respectively (**Fig. 4d**, white dashed line).

To further evaluate the performance of EmpirDE, we used the reproducibility of DEG calls to empirically evaluate the power and error of the DE analysis. We randomly selected and independently repeated, in triplicate, 36 RNAi experiments that yielded a broad range of DEGs (**Fig. 4e**). DEGs that were identified in one experiment but had no significant change in the other (FC < 1.1 or in reversed direction), are considered genuine false discoveries. We used the rate of such irreproducible DEGs to estimate the true FDR and found that EmpirDE showed a rate of irreproducible calls consistent with the FDR threshold (10%), regardless of the effect size (number of DEGs) (**Fig. 4f**). In contrast, DESeq2 analysis achieved the desired control of FDR only when the effect size was large. The rigorous control of false positives of the two-pronged EmpirDE approach can be further demonstrated by visually inspecting each of the 36 conditions (**Fig. 4g, h**, **Extended Data Fig. 4**). For instance, many DEGs in the *metr-1* RNAi experiment were control-outlier genes and, as expected, these did not replicate in the repeat experiment (**Fig. 4g**, right side, dashed black circles). Although non-control-outlier DEGs were generally reproduced with both DESeq2 and EmpirDE, the latter still eliminated a few highly changed but unreproduced calls (**Fig. 4g**, left). Notably, the EmpirDE approach was critical when true positives were sparse and false positives identified by DESeq2 analysis masked the retrieval of true positives (**Fig. 4h**). Finally, these analyses also facilitated EmpirDE parameter optimization, for instance, selecting an optimal threshold to determine control-outlier genes in the first step of the framework (**Supplementary Methods**, **Extended Data Fig. 3d**, e).

Taken together, with the EmpirDE framework that uses gene-specific empirical null models, WPS can robustly assign DEGs for large numbers of perturbations with high signal-to-noise ratio and rigorously controlled FDR for perturbations eliciting from a few to thousands of DEGs.

### A proof-of-principle of WPS with 103 NHRs

NHRs are a family of transcription factors (TFs) that play important roles in various physiological processes including metabolism, development, and homeostasis^47^. The *C. elegans* genome is predicted to encode more than 250 NHRs, making it the largest TF family. In contrast, the human genome encodes only 48^48–50^. Although many NHRs have been studied in *C. elegans*^21,51–55^, more than half remain completely uncharacterized^47,56^. We analyzed the expression levels and patterns of all 288 predicted NHRs encoded by the *C. elegans* genome and selected a set of 103 that are expressed at relatively high levels both in the whole body and in the intestine and/or hypodermis, tissues highly suitable for RNAi^57^, to test the WPS method (**Fig. 5a, b**).

**Fig. 5:**
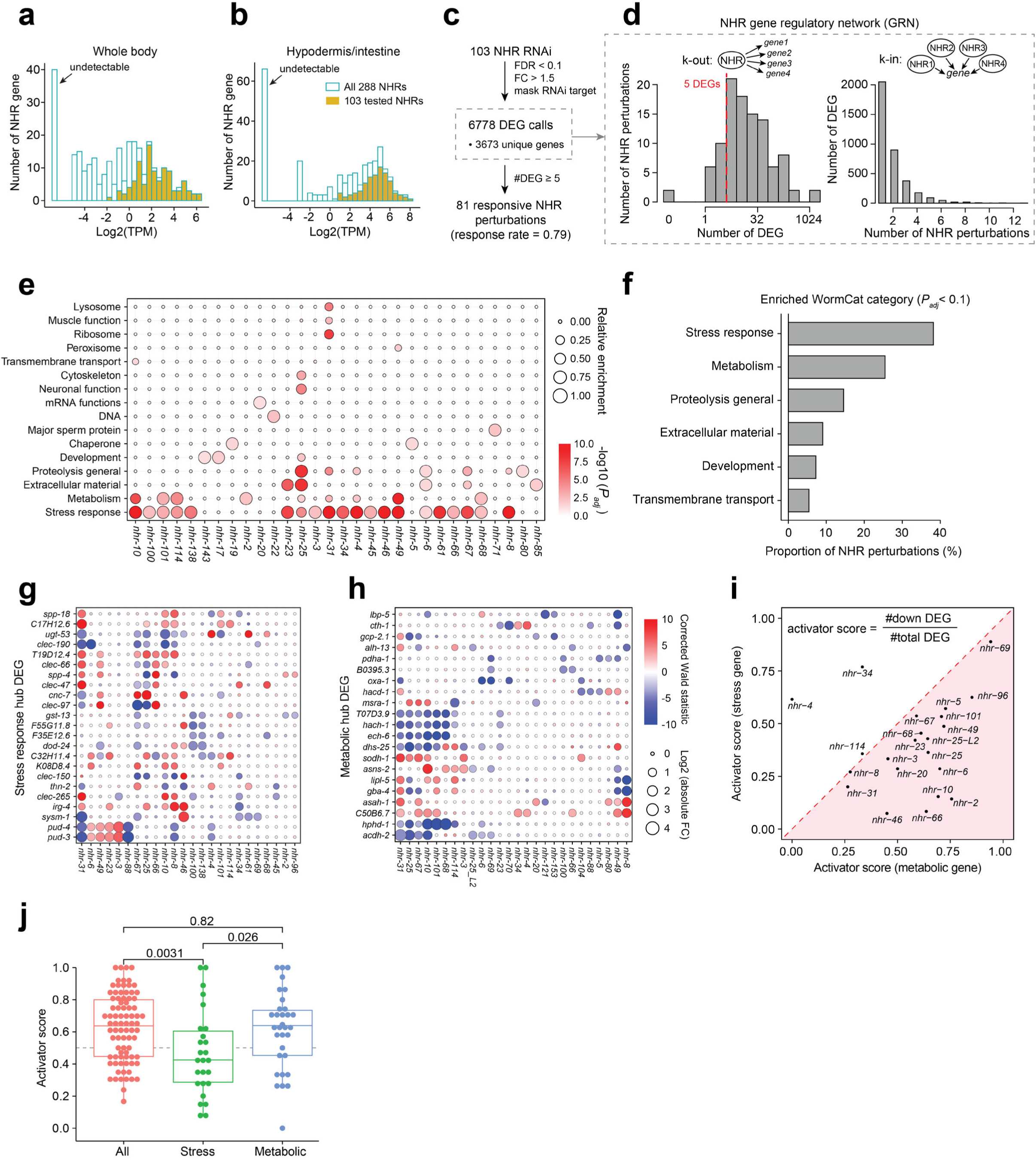
A NHR gene regulatory network. **a**, Expression levels of *nhr* genes in adult animals. The TPM was quantified by the reference profile used in **Extended Data Fig. 1c**. **b**, Tissue expression of *nhr* genes based on an adult stage single-cell RNA-seq data^71^. The maximal TPM of hypodermis and intestine expression is shown in the plot. **c**, Summary of NHR WPS experiments. **d**, Distribution of the number of DEGs in NHR perturbations (i.e., k-out) and the number of NHRs regulating the same gene (i.e., k-in). The red dashed line indicates the responsiveness threshold (≥ 5 DEGs). **e**, Functional enrichment analysis of NHR RNAi conditions. Only the 54 NHRs (55 perturbations because *nhr-25* was perturbed at both L1 and L2 stages) with more than 10 DEGs were analyzed to ensure power. Conditions without significant (*P_adj_* > 0.05) enrichment are not shown in the plot. The relative enrichment is defined as the -log10(*P_adj_*) normalized by its maximum in a given RNAi condition. **f**. Proportion of testable RNAi (i.e. > 10 DEGs) that showed significant enrichment (*P_adj_* < 0.1) in WormCat Level 1. **g**, **h**, Visualization of hub genes (i.e., regulated by >5 NHRs) of the stress response (**g**) and metabolic (**h**) categories. **i**, Comparison of the activator score for metabolic (x-axis) and stress response (y-axis) DEGs. NHR perturbations with ≥ 5 DEGs in the corresponding categories were analyzed. **j**, Overall comparison of activator scores for different DEG categories. Two-sample Wilcoxon tests were performed to calculate the *P* values.

WPS analysis of these NHRs yielded a gene regulatory network (GRN) comprising 6,778 interactions between 101 perturbations and 3,673 genes, with in- and out-degrees following expected distributions^58,59^ (**Fig. 5c, d**, **Supplementary Table 3**). We found that ∼80% of perturbed NHRs (81) were responsive (≥ 5 DEGs, a conservative threshold compared with the false positives in the NTP analysis, **Fig. 4d**). This rate is much higher than that reported in whole-genome single-cell Perturb-seq experiments (∼30%)^13^, and almost double than that we found in metabolic gene screens (40%) [REWIRING], indicating that the majority of the 103 NHRs tested are actively regulating gene expression in adult animals. Most NHR knockdowns resulted in moderate number of DEGs (5-100) (**Fig. 5d**) and the magnitude of gene expression changes were also modest (**Extended Data Fig. 5a**). Notably, 70% of DEGs identified in more than one perturbation changed in the same direction (up or down, **Extended Data Fig. 5b**). As expected, genes that were mostly down-regulated are expressed at higher levels than those that are mostly up-regulated in NHR perturbations (**Extended Data Fig. 5c**).

We used WormCat^60^ to identify biological processes enriched in the DEGs for each of the 54 NHRs that yielded more than 10 DEGs and found that many NHRs affected genes involved in stress response and metabolism, specifically pathogen response and lipid metabolism (**Fig. 5e, f** and **Extended Data Fig. 5d**, **e**). These observations indicate that several NHRs may function to establish and/or maintain metabolic functions and to prime the animal to respond to different stressors. The knockdown of individual NHRs was associated with several other WormCat categories as well. Some of these were known, including the association of *nhr-31* with the lysosomal vATPase, *nhr-49* with lipid metabolism, and *nhr*-*10, 68, 114, 101* with the propionate shunt pathway^54,61,62^ (**Extended Data Fig. 5d**, **f**).

We wondered whether the regulation of stress response and metabolic genes was due to a few hub genes that are affected by many NHR perturbations (**Extended Data Fig. 5g**). We identified a set of hub genes (in-degree > 5) that are annotated in WormCat as either stress response or metabolic genes and found that different NHRs influenced the expression of distinct stress and metabolic genes (**Fig. 5g, h**). Together with the in-degree distribution that shows that most genes are regulated only by few NHRs (**Fig. 5d**), this analysis indicates that the enrichment for stress response and metabolic genes in the NHR GRN is not driven by a few common genes. Interestingly, we found that the GRN mostly consists of activating interactions, i.e., upon knockdown of an NHR, gene levels tend to go down. However, on average, we found that stress response gene expression was increased upon NHR knockdown (**Fig. 5i, j**, **Extended Data Fig. 6 a-f**). These results indicate that NHRs ensure the correct active expression of metabolic genes, especially those involved in lipid metabolism, while they repress stress response genes.

Of the 288 *C. elegans* NHRs, at least 269 are from a substantial expansion and diversification of the HNF4 family^63^. This observation raises the question whether these expanded HNF4 family members regulate similar targets, or whether they evolved distinct and diverse functions. To answer this, we compared the target genes of the 81 NHRs in the GRN in more detail using perturbation-perturbation similarity and made a remarkable observation: these NHRs not only regulated various sets of targets, but also clustered into modules consisting of distinct NHR pairs, which we named ‘pairwise modularity’. In fact, 52 of 81 NHRs (64%) shared a significant overlap only with one other NHR (**Fig. 6a, b and Extended Data Fig. 7, 8a, b, Supplementary Table 4**), and such pairwise modularity cannot be observed by random (**Fig. 6c**, **Extended Data Fig. 7**). Our analysis identified numerous NHR pairs for which functional relationships were not yet known and which provide hypotheses for further experimental evaluation (two examples in **Extended Data Fig. 8c**, **d**). Notably, this observation was facilitated by the EmpirDE framework because it increased the signal-to-noise ratio compared to DESeq2 (**Extended Data Fig. 7**). Thus, even with a relatively low number of perturbations (∼100 RNAi conditions), EmpirDE can effectively increase the interpretability of the data.

**Fig. 6:**
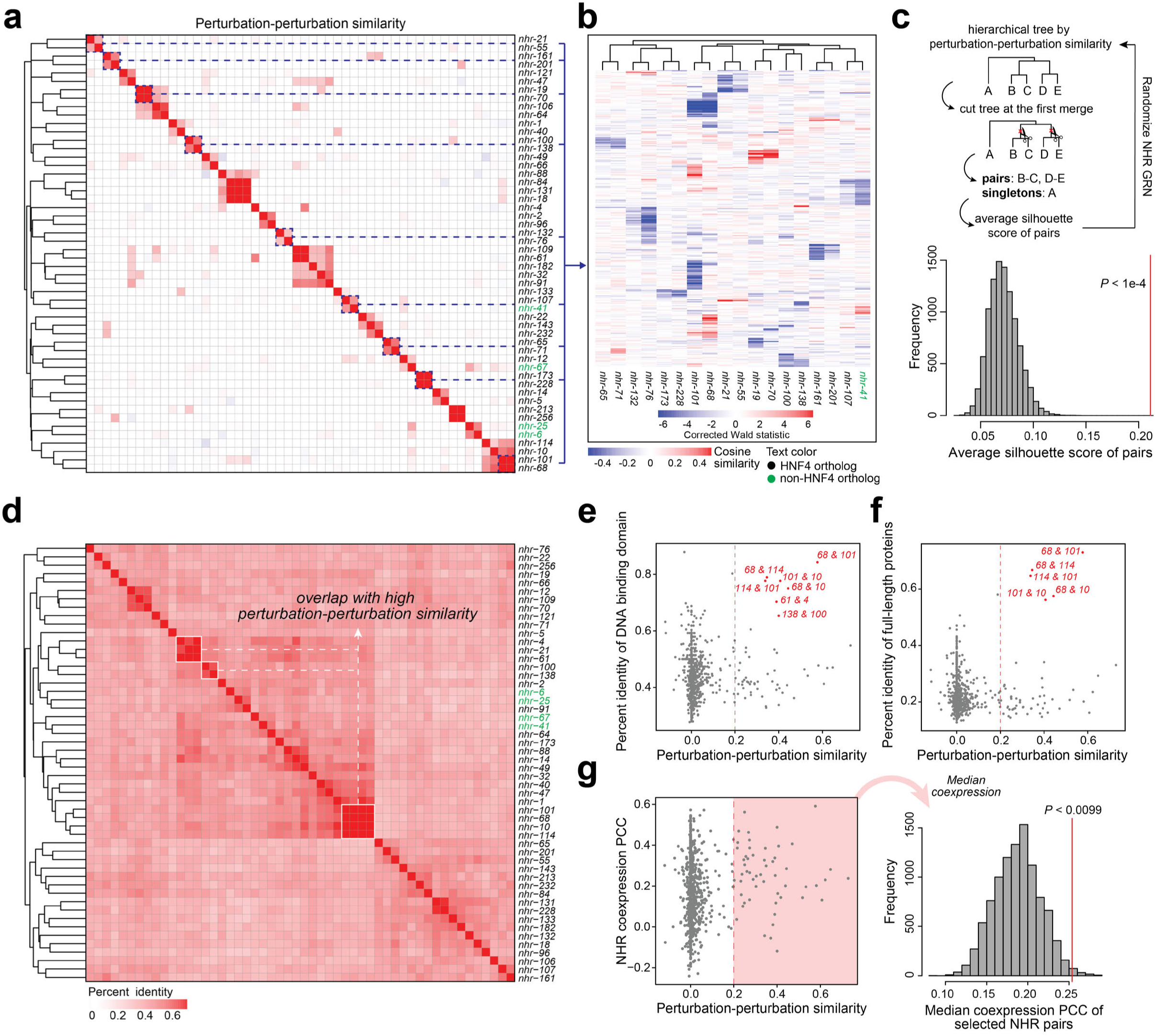
The pairwise modularity of NHRs. **a**, Heatmap depicting perturbation-perturbation similarity of DEG profiles for the NHR perturbations. The perturbation-perturbation similarity was defined by cosine similarity of the filtered log2(FC) profile. The filtered log2(FC) was derived by masking the log2(FC) values of genes that are not called as DEGs (FDR < 0.1, FC > 1.5) to zero. **b**, Visualization of gene expression changes in selected NHR pairs. The gene expression change was measured by the corrected Wald statistic. Rows are the union DEGs of these selected NHR perturbations. **c**, Randomization test of the pairwise modularity of NHR gene family. The schematic shows the design of the randomization test. Histogram shows the average silhouette score of pairs in 10,000 randomizations. The red line indicates the observed score from real data. **d**, Heatmap depicting protein sequence similarity (percent identity) for the DNA binding domain (DBD) of NHRs. The heatmap was clustered using distance matrix generated by Clustal Omega online tool from EMBL-EBI^72^ (**Supplementary Methods**). **e, f**, Scatter plots showing the comparison between perturbation-perturbation similarity and sequence similarity of DBD (**e**) and full-length protein (**f**). Each data point indicates a pair of NHRs and selected pairs are labeled. **g**, Scatter plot and randomization test for the associations between perturbation-perturbation similarity and NHR coexpression. Coexpression was measured based on the median Pearson Correlation Coefficient (PCC) of *nhr* gene expression in a compendium of *C. elegans* gene expression data across various conditions^73^ (**Supplementary Methods**). The median coexpression level of pairs with a cosine similarity greater than 0.2 (red region in (**g**)) is calculated and compared with that from randomized data (**Supplementary Methods**). The histogram shows that the median from real data (red line) is significantly greater than that from randomized data, indicating a statistically significant association between NHR coexpression and perturbation-perturbation similarity.

Interestingly, NHR sequence similarity only correlated with few pairs that shared target genes. Overall, protein sequences of the NHR DNA binding domains showed relative low similarity to each other (percent identity < 0.5), with only limited number of clusters (**Fig. 6d**). Although the few pairs with high protein sequence similarity were more likely to share targets (e.g., *nhr-10, nhr-68, nhr-114,* and *nhr-101*) the protein similarity between most NHR pairs was lower and did not correlate with perturbation-perturbation similarity (**Fig. 6e, f**). Remarkably, even some NHRs from different evolutionary origins significantly shared targets (e.g., *nhr-107* and *nhr-41,* **Fig. 6a**). In comparison, we found that perturbation-perturbation similarity correlated better with *nhr* coexpression levels (**Fig. 6g**). Therefore, the pairwise modularity unveiled by WPS may be affected more by mechanisms involved in the regulation, and less by the biophysical properties (e.g., DNA binding domains), of these NHRs.

## Discussion

In this study, we provide a WPS platform that combines strengths of single-cell and bulk RNA sequencing with whole animal gene perturbations by RNAi. WPS is both efficient and cost-effective, *e.g.,* a 2-week timeframe for collecting 96 perturbations in triplicate and more than 10-fold cost reduction compared with conventional methods (**Supplementary Protocols**), which enables replicate screens with full transcriptome readouts of hundreds of perturbations in a living animal. Future screens with additional RNAi libraries, different bacterial diets, and supplementation of metabolites or drugs will provide insights into how the animal responds to a variety of perturbations.

A key advantage of WPS is that it is based on whole-organism *in vivo* perturbations. While this is not feasible in mammals, it should be applicable to organisms amenable to large-scale RNAi screens, such as Drosophila. However, we do envision that WPS-like screens will be feasible in bulk in tissue culture cells, especially when smaller sub-libraries of genes (∼100) are selected for perturbations. Another key feature of WPS is the EmpirDE framework that uses an empirical null for each detected gene and that can be applied due to the scale of WPS experiments, and which can alleviate systematic errors such as batch effects. Our study also shows and corrects the anti-conservative *P* values that result from incorrect theoretical assumptions of parametric differential gene expression models, which is in line with recent studies^39,40^. Although the concept of empirical null has been widely applied in genomics^64–66^, to the best of our knowledge, it has not been used to directly model the test statistic distribution at the level of individual genes (features), possibly due to the lack of such massively parallel data generated with WPS. EmpirDE exploits the unique power of having many conditions (> 100), each with three replicates, to achieve rigorous statistical analyses. Conventionally, such level of statistical rigor is only achievable with a high number of replicates (e.g., 8-12)^67^, which is a rare occasion in biological research. Given the dramatic decrease in cost and increase in throughput for RNA-seq in recent years ^44,68,69^, we envision the soon explosion of large-scale RNA-seq datasets in both research and clinic settings, all of which will benefit from using EmpirDE.

By applying WPS to more than 100 NHR perturbations, we discover a pairwise modularity in which two or more NHRs regulate the expression of overlapping sets of genes, which cannot be explained by protein (and presumably binding site) similarity. Instead, this pairwise modularity suggests that ‘AND-logic gates’ are a common mechanism of gene regulation in *C. elegans*. Future studies with other TFs will be important to see if this is a general principle, or if it is a specific feature of NHRs. Knockdown of many NHRs affected only few genes, suggesting that these TFs may either not be active under the conditions tested, or are truly specialized in their regulatory function. In two companion studies [WIRING][REWIRING], we further validated WPS by perturbating ∼900 metabolic genes. These studies generated high-quality, highly interpretable datasets, providing tremendous insights into metabolic wiring and rewiring at a systems level. Notably, WPS interrogates gene functions *in vivo*, thus linking genes to their native physiological roles. For instance, using metabolic gene WPS data, we identified an unconventional central carbon metabolism that consumes ribose, rather than glucose, from dietary RNA and through the pentose phosphate pathway. Together, we envision that WPS-style *in vivo* functional genomics will provide a powerful tool to uncover gene functions in living organisms.

## SUPPLEMENTARY TABLES LEGENDS

Supplementary Table 1: The reference TPM profile of adult *C. elegans* used in this study. This is obtained from the combined vector control data collected at 62 to 65 hours post L1 stage in one biological replicate (See Supplementary Methods for details).

Supplementary Table 2: List of 67 non-targeting perturbations (NTPs). The list does not include the 4 random spike-in vector controls.

Supplementary Table 3: Differentially expressed genes in NHR perturbations.

Supplementary Table 4: Perturbation-perturbation similarity matrix of NHR perturbations (only includes responsive conditions).

## METHODS

See Supplementary Information.

## ACKNOWLEDGEMENTS

We thank members of the Walhout lab, Mike Lee, and Chad Myers for discussion and critical reading of the manuscript. We thank former Garber lab members, Kyle Gellatly and Rachel Murphy, for their help in the early stage of this project. This work was supported by grants from the National Institutes of Health R35GM122507 and DK068429 to A.J.M.W., U01HG012064 to M.G., and NSF DBI-1846216 and NIGMS R35GM140888 to J.J.L.

## AUTHOR CONTRIBUTIONS

H.Z., X.L., and A.J.M.W. conceived the project and wrote the manuscript. H.Z. and X.L. jointly developed the WPS technology and analyzed the data. H.Z. conducted the experiments. X.L. wrote the codes. D.S. helped design the simulation study under the supervision of J.J.L. D.S. also provided critical discussions that led to the idea of modeling the empirical null. O.Y. and A.K. helped with the dolphinNext pipeline for WPS data processing. S.N. provided the NHR coexpression data. M.G. and A.J.M.W. supervised the study. The co-first authorship order was determined by a coin flip. XL and HZ contributed equally and reserve the right to list their name first in their resumes.

## ADDITIONAL INFORMATION

Correspondence and requests for materials should be addressed to A.J.M.W., or M.G. Supplementary Information is available for this paper.

## DATA AVAILABILITY

The WPS data analysis pipeline is available at https://github.com/XuhangLi/WPS including detailed procedures of raw data processing, quality control and EmpirDE analysis. A walkthrough of the pipeline can be found in the repository. A standalone R package for the EmpirDE analysis can be found in https://github.com/XuhangLi/EmpirDE. All other source code to reproduce the figures and analyses in this study will be deposited to Zenodo before publication. Raw and processed data in this study are available in Gene Expression Omnibus (GEO) session GSE255865. Downloadable read count data are also available at the WPS portal hosted in our WormFlux website https://wormflux.umassmed.edu/WPS.

**Extended Data Fig. 1:**
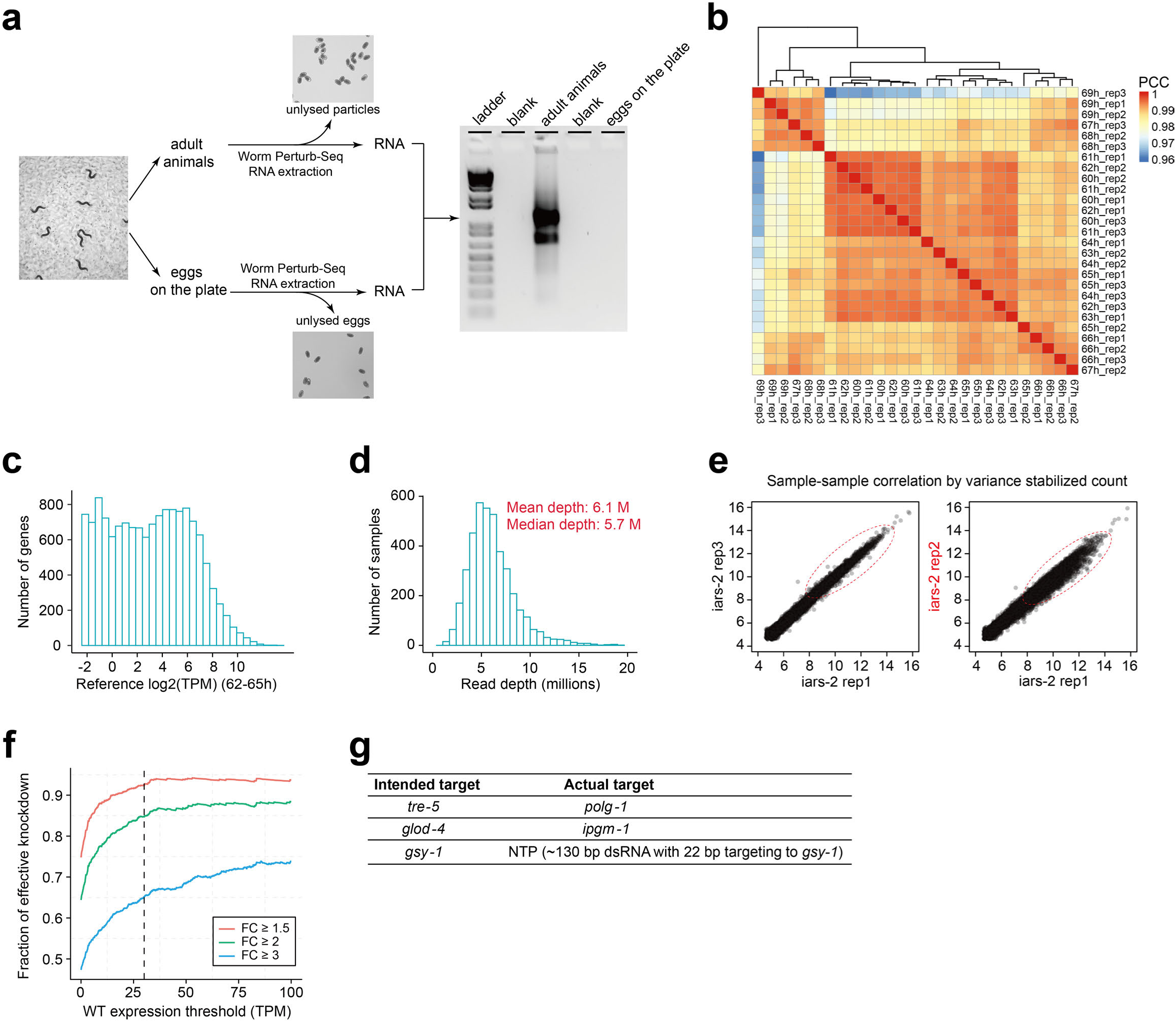
Development of WPS. **a**, WPS RNA extraction method does not extract RNA from eggs. **b**, Zoom-in heatmap of Fig. 2a. **c**, Histogram of gene expression in the subsampling reference profile. The reference was generated by combining the data of a single replicate from 62 to 65 hours in (**b**). **d**, Read depth distribution of 4,083 WPS samples combining metabolic and NHR WPS datasets. **e**, Examples of gene expression correlation between replicates with good (left) and poor (right, labeled in red) quality. **f**, Fractions of perturbations with effective targeted gene knockdown versus targeted gene expression. Effective knockdown is defined by a minimum decrease of 1.5 (red), 2 (green) and 3 (blue) folds in the targeted gene expression, based on differential expression analysis. The fraction was calculated using 1,048 valid RNAi conditions from both metabolic and NHR WPS datasets, titrated with a minimum targeted gene expression threshold indicated in x-axis. The Wild-Type (WT) expression was based on the reference profile in (**c**). **g**, Sanger sequencing results of the three fail-QC conditions in Fig. 2f.

**Extended Data Fig. 2:**
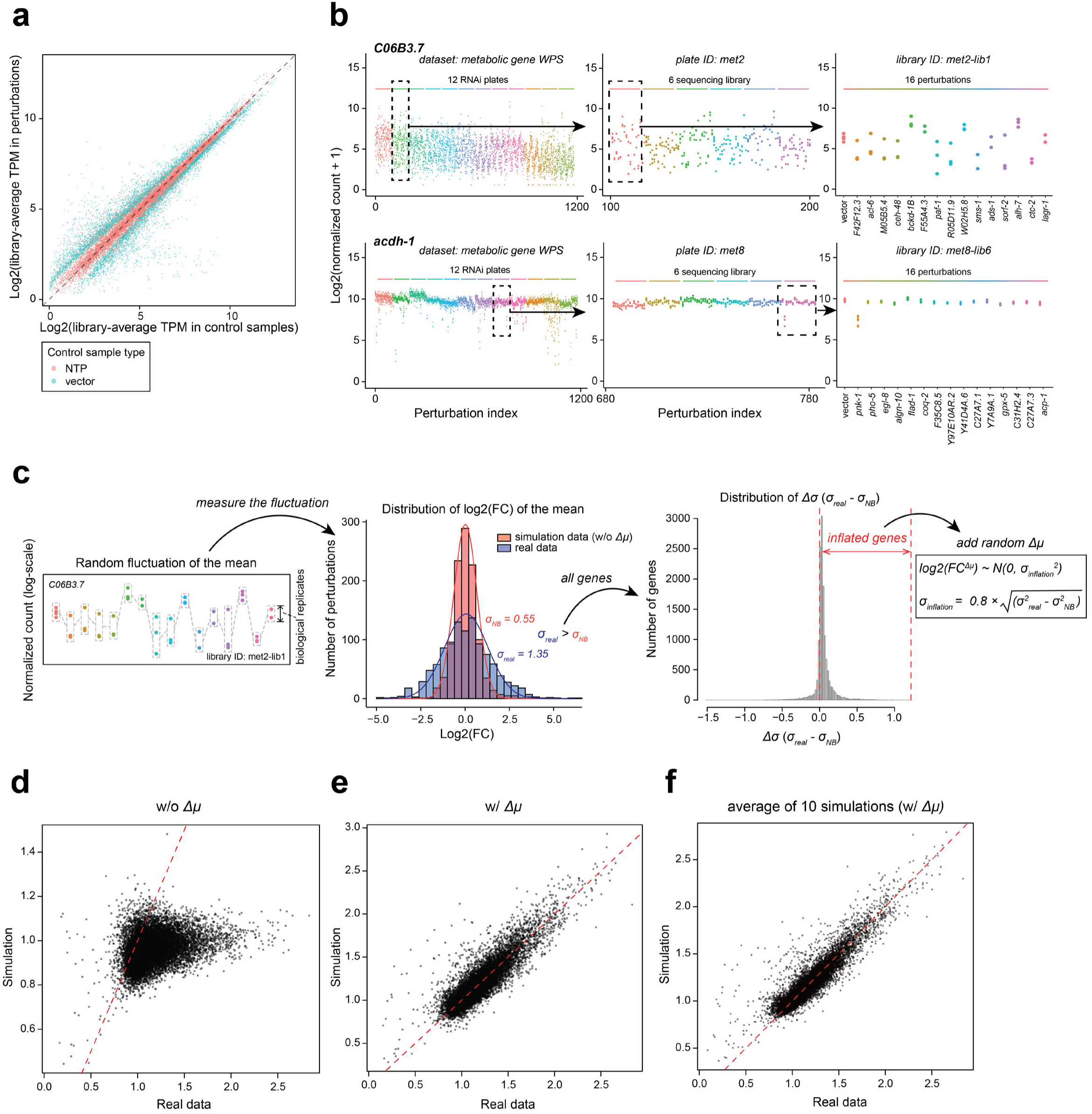
Investigating the source of false discovery in WPS analysis. **a**, Scatter plot showing the expression level of control-outlier genes in different types of samples. The control-outlier genes in the 72 metabolic-gene WPS sequencing libraries are combined and displayed in the plot and each data point refers to a control-outlier gene in a sequencing library. The y-axis represents the average TPM of all RNAi samples (excluding NTPs and vector controls) in a sequencing library, which is compared to the average TPM of either NTP (red) or vector control (blue) samples in the same sequencing library on the x-axis. **b**, Gene expression levels of two example genes exhibiting high (top) or moderate (down) fluctuation of the mean. The three panels from left to right present a sequential zoom-in view of the expression pattern at the level of entire metabolic-gene WPS dataset, an RNAi plate, and a sequencing library. The perturbation index is an arbitrary index indicating the number of perturbations displayed in the plot. **c**, Illustration of the quantification and simulation of random fluctuation of the mean. To quantify the fluctuation level of each gene, a distribution of its log2(FC) across the thousand RNAi conditions in the metabolic-gene WPS dataset was fitted by the same procedure used for fitting the empirical null of the Wald statistic. The fitted sigma from real data was compared to that from the simulation data without *Δμ*. Inflation of the sigma level compared with simulation fit was used as a proxy of mean fluctuation to create a heuristic (formula on the right) for generating gene-specific random *Δμ* (**Supplementary Methods**). **d**, **e**, Comparison of the fitted standard deviation of the Wald statistic between simulation and real data for all genes. The red dashed line indicates the diagonal (y=x). The scatter plot shows results from one simulation of the metabolic-gene WPS dataset. **f**, Similar to (**e**), but showing the average of fitted standard deviations in 10 independent simulations.

**Extended Data Fig. 3:**
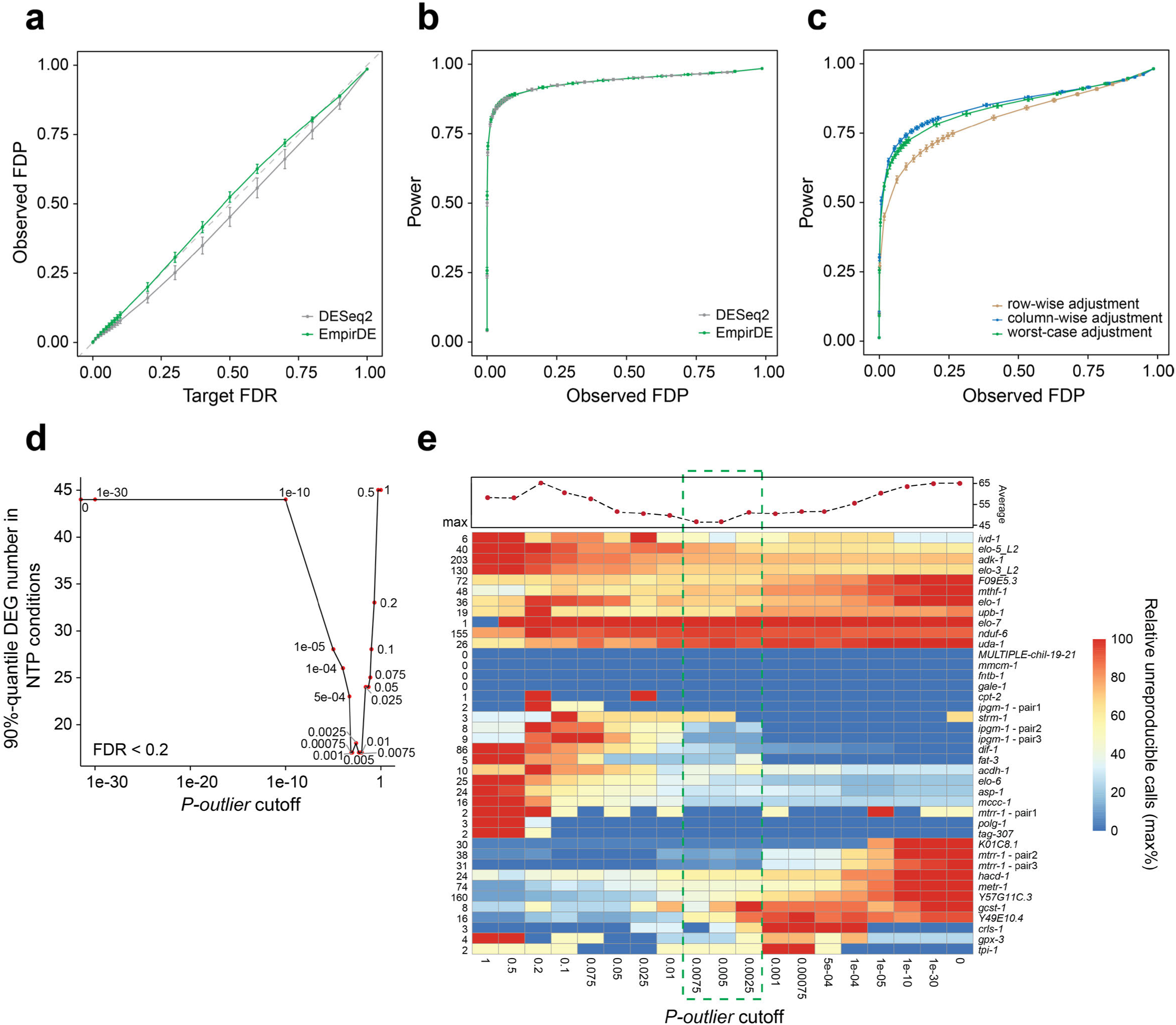
Benchmarking and parameter optimization of EmpirDE analysis framework. **a**, **b**, Performance benchmarking of the EmpirDE analysis using simulation data without *Δμ.* FDP, false discovery proportion. **c**, Benchmarking the FDR control of different multiple testing adjustment strategies. **d, e,** Two evaluations of false discovery to select the threshold for defining control-outlier genes in the first step of EmpirDE analysis. (**d**) shows the false discovery estimation by DEG calls in NTP samples across different *P*-outlier thresholds, and (**e**) shows the estimation by unreproducible calls in the 36 repeated conditions. To increase the sensitivity of the analysis, DEGs in (**d**) were solely defined with FDR threshold of 0.2. Colors in (**e**) show the fraction of all unreproducible DEGs with respect to the maximum level in each condition across a range of *P-outlier* threshold. The maximum levels of unreproducible DEGs are shown on the left of the heatmap. Scatter plot on the top shows the column-wise average (only conditions with maximum level greater than 5 DEGs were used for better robustness). For definition of unreproducible DEGs, please refer to **Supplementary Methods**.

**Extended Data Fig. 4:**
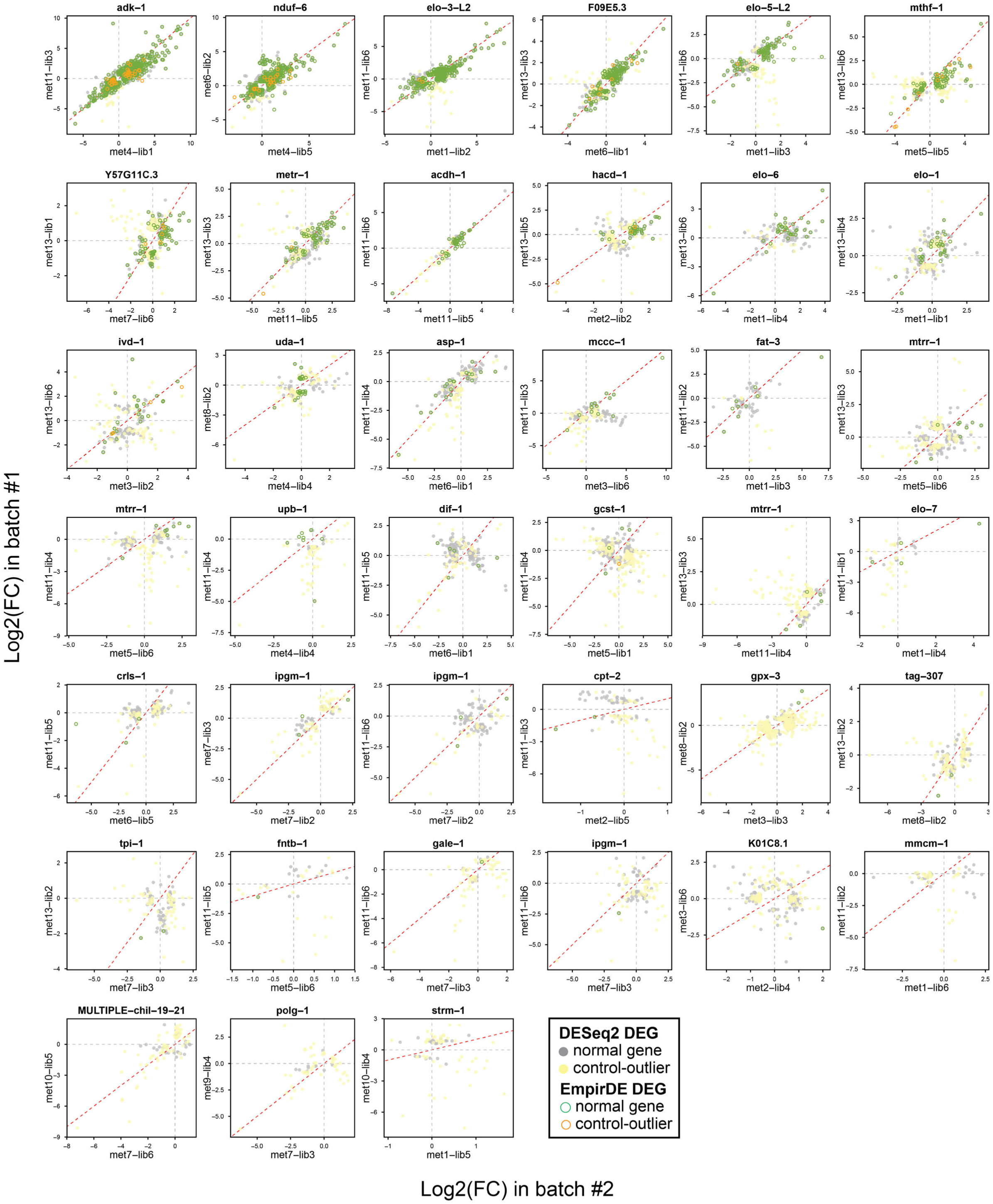
WPS analysis reproducibility. Thirty-six WPS experiments were repeated, and some were done more than twice (e.g., *mtrr-1*). Each experiment included triplicate perturbations. Plots show comparison of the log2(FC) of genes called as DEGs in either experiment. Red dashed lines indicate the diagonal (y=x).

**Extended Data Fig. 5:**
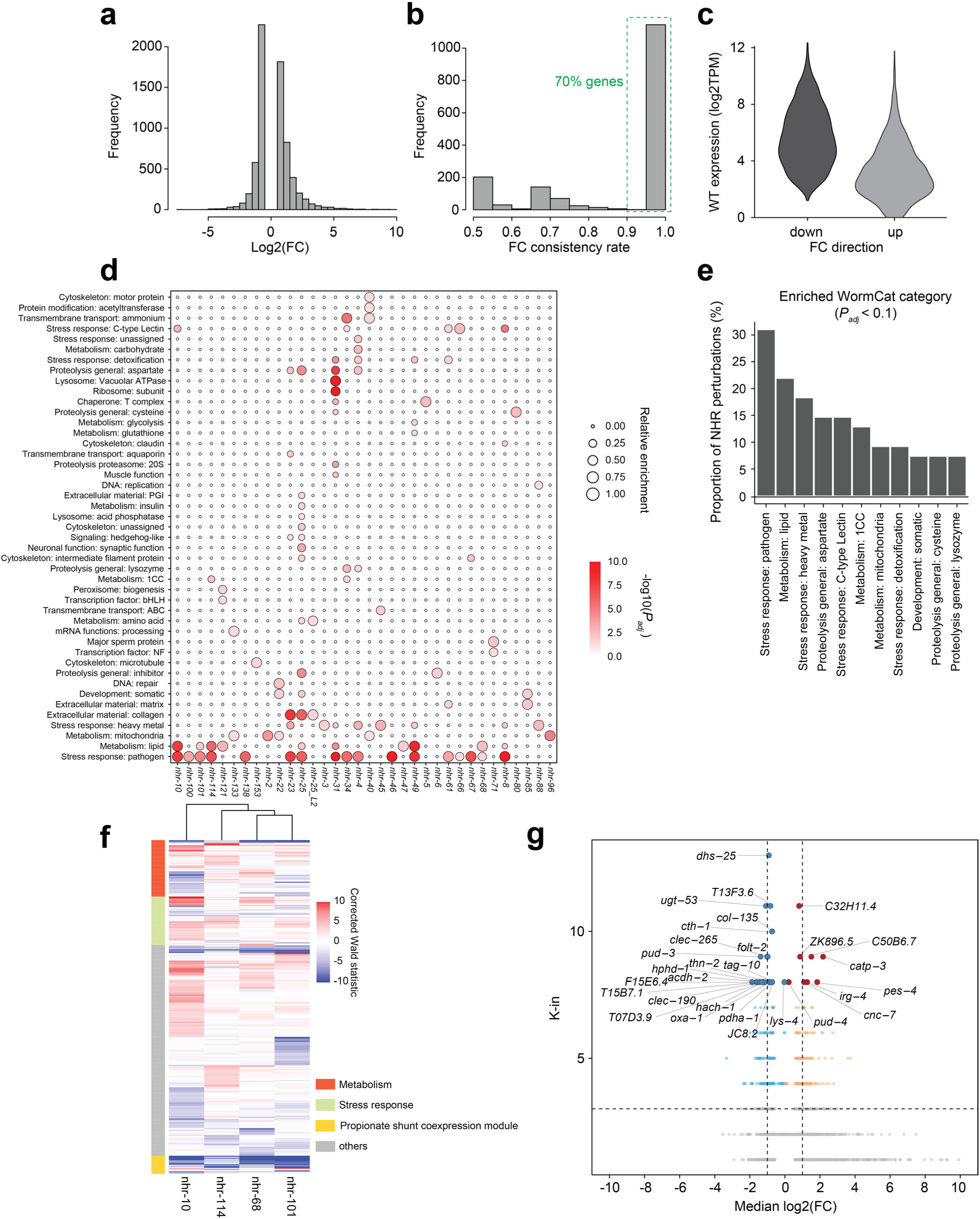
Analysis of NHR WPS data. **a**, Log2(FC) distribution of all DEGs in the NHR WPS experiments. **b**, Distribution of the fold change consistency (up or down) for all DEGs with k-in ≥ 2, which was defined as the proportion of DEG calls that increase or decrease in the WPS experiments (whichever greater). **c**, Association between fold change direction and basal expression level of the DEGs. **d**, Gene set enrichment analysis of NHR perturbations using WormCat Level 2 categories^60^. Only NHR perturbations with more than 10 DEG calls were analyzed. Conditions without significant (*P_adj_* > 0.05) enrichment are not shown in the plot. The relative enrichment is defined as the -log10(*P_adj_*) normalized by its maximum in a given perturbation. **e,** Proportion of testable perturbations that showed significant enrichment (*P_adj_* < 0.1) in WormCat Levels 2 categories. **f**, Heatmap displaying the corrected Wald statistics for the union of DEGs in *nhr-10*, *nhr-68*, *nhr-101*, and *nhr-114* perturbations. The propionate shunt coexpression module was defined based on a previous study^73^. **g,** Volcano plot showing the hub DEGs in the NHR GRN. The median log2(FC) is the median level of the log2(FC) values of all interactions for a corresponding gene in the GRN. The red and blue colors are used to indicate up and down regulated hub DEGs, respectively.

**Extended Data Fig. 6:**
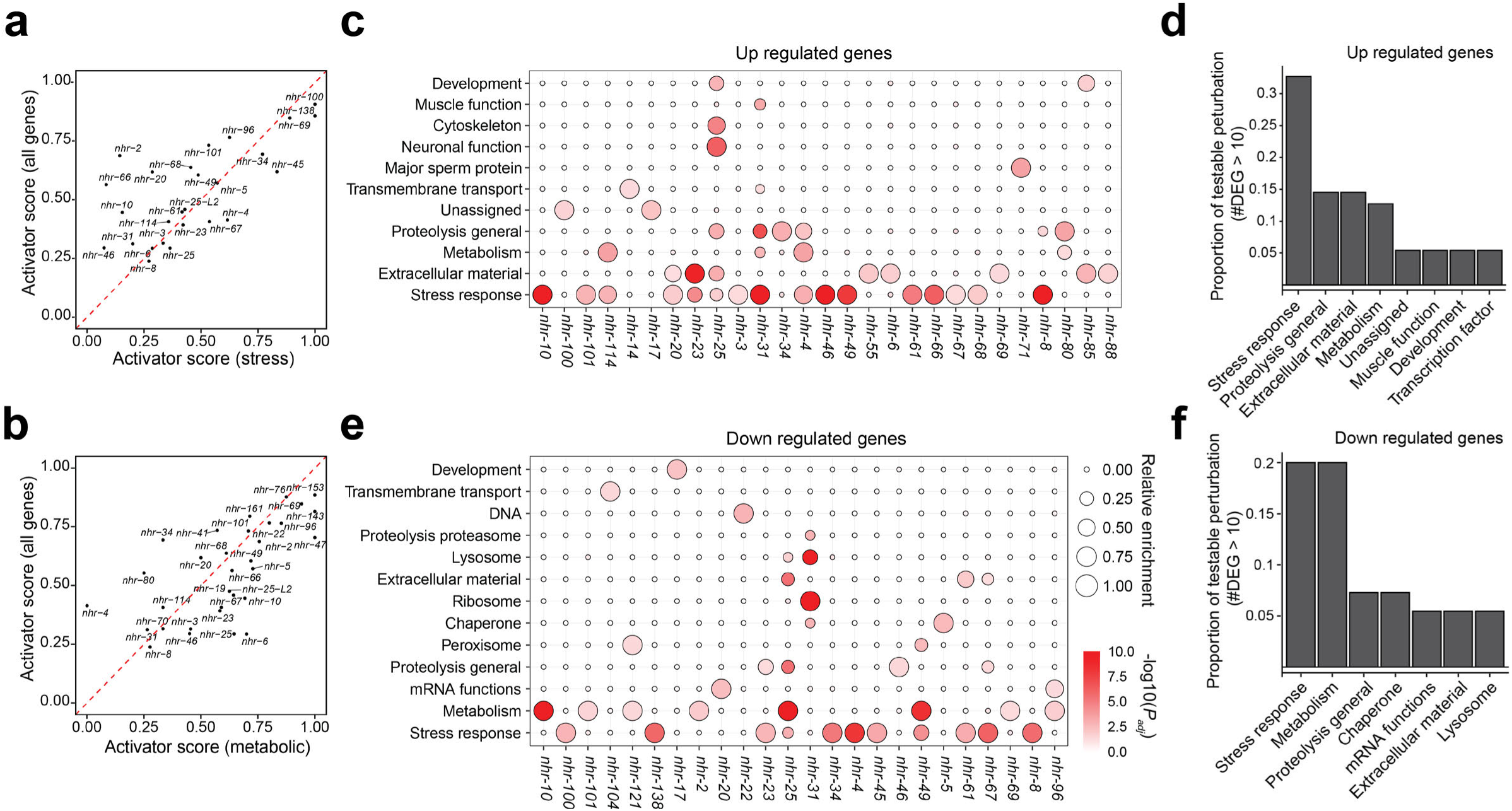
Analysis of NHR GRN. **a**, **b**, Correlation between the activator score for all DEGs and that for stress response (**a**) and metabolic (**b**) genes. The stress response and metabolic genes were defined based on WormCat Level 1 categories^60^. Only perturbations with ≥ 5 stress (**a**) or metabolic (**b**) DEGs are shown in the plot. **c, d**, **e**, **f**, Gene set enrichment analysis of up-regulated (**c**,**d**) and down-regulated (**e**,**f**) genes. WormCat Level 1 categories were used as the gene sets.

**Extended Data Fig. 7:**
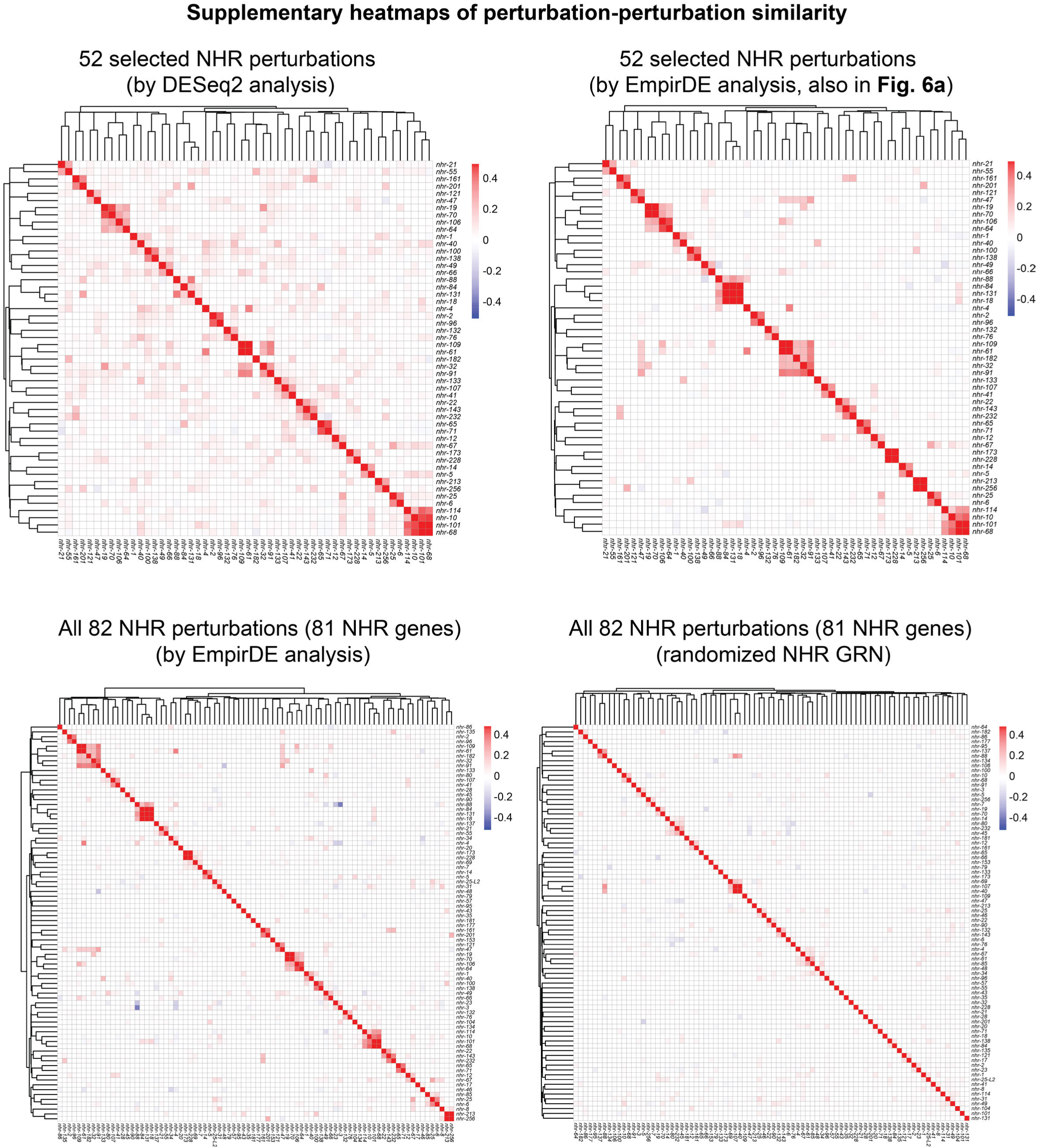
Supplementary heatmaps of perturbation-perturbation similarity.

**Extended Data Fig. 8:**
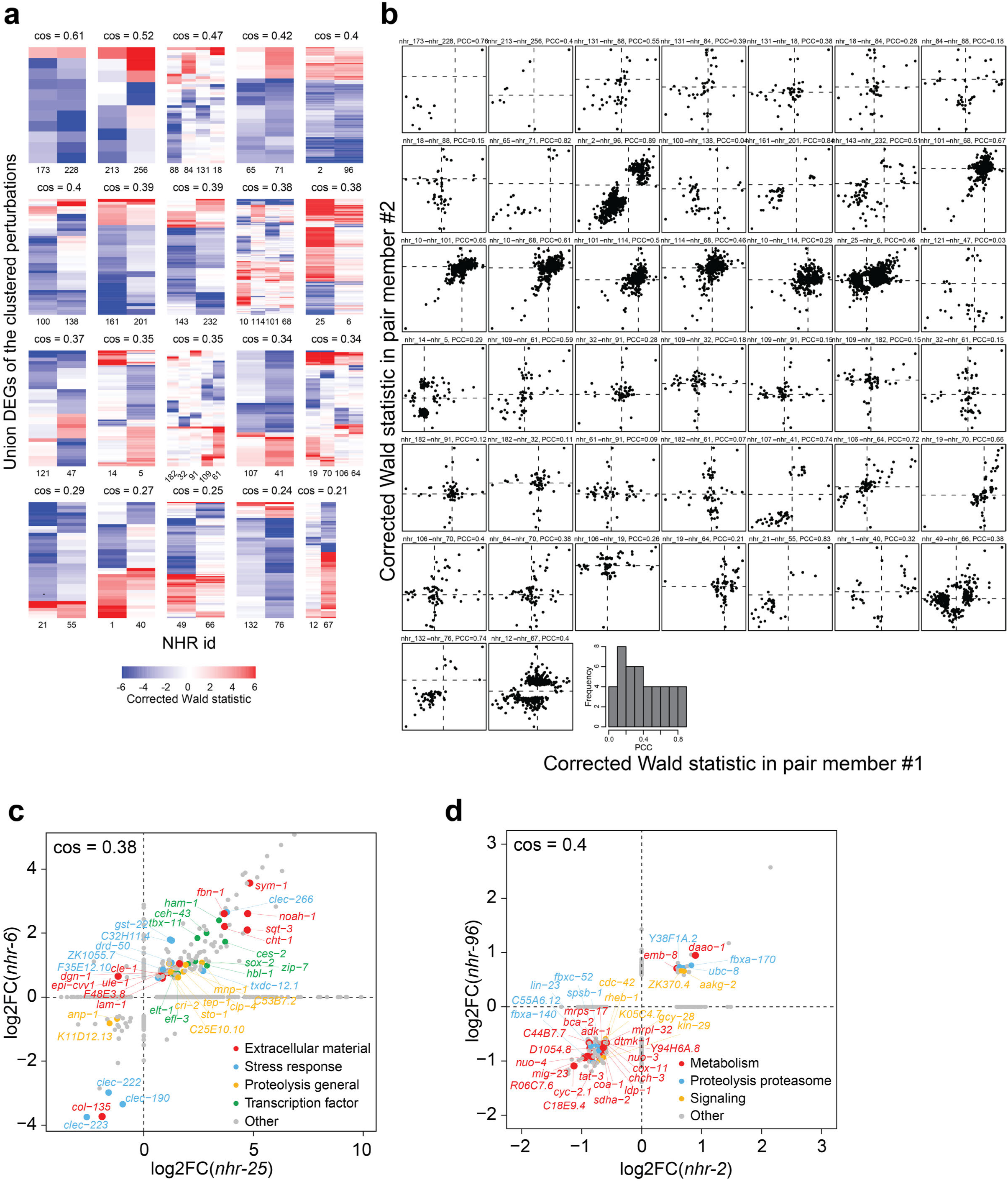
Visualization of the gene expression changes of selected NHR perturbation clusters. **a**, Heatmaps showing the corrected Wald statistic. The NHR perturbations were grouped by clusters. These clusters were identified by cutting the hierarchical tree in Fig. 6a using *cutree* function in R with h = 0.9. The union DEGs of conditions in each cluster are shown in the y-axis. The numbers in x-axis indicate the number in the corresponding *nhr* gene name. cos, cosine similarity. When there are more than two conditions in a cluster, the average pairwise cosine similarity was labeled. **b**, Scatter plots showing the correlation of expression changes in NHR pairs. The NHR pairs were defined as the intra-cluster pairs in (**a**) and the corrected Wald statistics for the union DEGs of the pair were shown in the plot. A histogram for the Pearson Correlation Coefficient (PCC) of each pair is shown at the bottom. **c**, **d**, Comparison of filtered log2(FC) profile for two example pairs of NHR perturbations. The union set of DEGs in the two conditions are shown in the plot. cos, cosine similarity.

