## Supplementary Information for "Worm Perturb-Seq: massively parallel whole-animal RNAi and RNA-seq"

**This PDF file includes:**

- Supplementary Notes
- Supplementary Methods
- Supplementary Protocols

**Other Supplementary Information for this manuscript include the following:**

- Supplementary Table 1 to 4

### Supplementary Notes

Supplementary Note 1: Overview of RNAi bacterial library preparation, RNA extraction and multiplexed sequencing library construction for Worm Perturb-Seq (WPS)

To start an WPS experiment, an RNAi bacterial library was prepared for genes of interest, each from a single colony of bacteria harboring the corresponding RNAi construct. In each WPS experiment, RNAi bacteria were grown to log phase before being seeded in 6-well solid-media plates. Bacterial expression of double stranded RNA for RNAi was induced by Isopropyl β-D-1-thiogalactopyranoside (IPTG) and ~200 *C. elegans* at the first larval stage (L1) were added to each well. When grown to the desired stage, animals were harvested in a 96-well plate for RNA extraction.

We developed a protocol for the high-throughput RNA extraction in 96-well plates (**Supplementary Protocol**). The most frequently used methods for RNA extraction use Trizol and multiple liquid nitrogen freeze/thaw cycles to break the animal's cuticle in individual tubes, which does not enable easy high-throughput experiments. Therefore, we explored whether we could use a combination of SDS and DTT, which has previously been used to break the animal's cuticle[1]. We used this method in combination with an optimized chemical cocktail containing proteinase K, commonly used worm lysis buffer[2], and a commercially available tissue lysing reagent to lyse the animal in 96-well plates. Importantly, this approach does not lyse eggs, making it suitable to use WPS for gravid adults (**Extended Data Fig. 1a**). After lysis, RNA was extracted using 96-well silica columns[3] (**Supplementary Protocol**). Finally, RNA concentrations were normalized to facilitate even pooling of samples in the next step.

To generate multiplexed RNA-seq libraries, we optimized the 3' end barcoding method CEL-Seq2[4], which was originally developed for single-cell RNA-seq[5]. We optimized the protocol for bulk RNA-seq application by reducing the use of costly reagents, such as reverse transcriptase, and found that a general 10x reduction produced libraries with similar quality and yield. We also optimized other experimental details such as incubation times, substrate concentrations, Quality Control (QC) checkpoints, and an earlier pooling of samples (**Fig. 1b** and **Supplementary Protocol**).

Supplementary Note 2: control-outlier genes and control-independent DE analysis

In the first step of EmpirDE, we performed Differentially Expression (DE) analysis at the level of individual sequencing libraries that consisted of ~16 conditions in triplicates (**Fig. 3d**, top). We first identified control-outlier genes as those consistently up- or down-regulated in at least three-quarters of conditions in a sequencing library, using a *P* value threshold (see **Supplementary Methods**). We found that most sequencing libraries contained fewer than 200 control-outlier genes (**Fig. 3e**). Importantly, albeit being consistently changed when compared with vector controls, the expression levels of these control-outlier genes in RNAi conditions were not systematically shifted when compared with the NTP condition(s) in the same sequencing library (**Extended Data Fig. 2a**). This validated that it was the gene expression in the vector control samples that was confounded. For these genes, we developed a control-independent DE analysis (**Fig. 3d,** top, algorithmic details are in **Supplementary Methods**). Briefly, for each control-outlier gene, we empirically identified a control-independent null population based on the distribution of its expression levels in a sequencing library. We applied an algorithm used by the GTEx consortium[6] to fit the distribution and to distinguish the main population that includes most conditions from outliers. Assuming the gene expression is not affected by most perturbations, this main population of the gene expression levels can be used as a null (**Fig. 3d**). We then performed DESeq2 by comparing the level for the gene in each RNAi condition to the newly defined null level. If a gene frequently appears as a control-outlier in different sequencing libraries (*i.e.,* > 25% of the libraries), we performed control-independent DE analysis for this gene in all libraries. For most regular, non-control-outlier genes, we used regular DESeq2 analysis, *i.e.,* compared each RNAi condition with the vector control from the same sequencing library to calculate the DE statistics (**Fig. 3d**).

### Supplementary Methods

#### *C. elegans* strains and maintenance

N2 strain was used as the wild-type strain. Animals were maintained at 20°C on solid nematode growth media (NGM)[7] and fed *E. coli* HT115 containing the empty RNAi vector L4440[8].

#### RNA interference

To construct WPS RNAi libraries, each RNAi bacteria strain was cherry picked from a parent library (e.g., the metabolic RNAi library[9], or TF RNAi library[10]) and streaked onto LB agar plate with 50 µg/mL ampicillin to produce single colonies. A single colony for each RNAi was used in the WPS RNAi library. RNAi was performed accordingly as described with slight modifications[11]. Briefly, bacteria were cultured overnight at 37°C in 1 mL LB supplemented with 50 µg/mL ampicillin in a 96-deep well plate. 100 µL of each culture was then diluted 50-fold using fresh LB medium with 50 µg/mL ampicillin in a well of a 24-deep well plate. After incubating for 4 hours at 37°C, bacteria were centrifuged at 3,000 g for 20 minutes in a Beckman Coulter Avanti® J-26XP High-Performance Centrifuge with a JS-5.3 Swing Bucket Centrifuge Rotor, and the pellet was resuspended in 200 µL M9. The resuspended bacteria were then transferred to 6-well NGM plates containing 50 µg/mL ampicillin and 2 mM Isopropyl β-d-1-thiogalactopyranoside (IPTG, Fisher Scientific) for induction of double-stranded RNA (dsRNA) expression. Plates were dried in a hood and incubated overnight at room temperature.

For developmental stage time course experiments, all animals were fed bacteria with vector control RNAi. Approximately 2,500 synchronized L1 animals were plated for collecting L2 animals; ~1000 synchronized L1 animals were plated for collecting L3 animals; ~500 synchronized L1 animals were plated for collecting L4 animals; and ~200 synchronized L1 animals were plated for young adult and gravid adult samples.

For all other WPS experiments, approximately 200 synchronized L1 animals were plated into each well, followed by incubation at 20°C for approximately 63 hours. In the cases where RNAi feeding led to a developmental delay phenotype, synchronized L1 animals were initially fed with vector control RNAi. After a period of 17 hours post-plating (i.e., at the L2 stage) or 25 hours post-plating (i.e., at the L3 stage), animals were transferred to the corresponding RNAi plates to circumvent RNAi-associated developmental delay. Animals usually develop normally after such delayed RNAi exposure. Only RNAi conditions without notable developmental delay were sequenced.

#### RNA extraction in 96-well plate

We developed a 96-well RNA extraction method for *C. elegans* tissues while leaving all eggs intact. Please refer to **Supplementary Protocol** for details.

#### WPS sequencing library construction

We adapted the CEL-Seq2 single-cell RNA-seq library construction protocol[5] for WPS sequencing library construction. We meticulously optimized each step of the protocol to ensure robustness and reproducibility. As part of this optimization, we modified the adaptor sequences of the CEL-Seq2 primers to ensure compatibility with both Illumina and BGI platforms for sequencing. For a comprehensive description of the modified protocol and primer sequences, please refer to the **Supplementary Protocol**.

#### WPS sequencing library design

We used an Illumina NextSeq sequencer capable of providing ~350 million reads (or its equivalent from BGI), which allowed us to pool ~50 samples to obtain an average coverage of ~7 million raw reads/sample. Typically, a library includes 15-16 RNAi conditions in triplicate and 6 vector control samples. We conducted three biological replicates for all RNAi conditions on different days. In each different-day replication, we included two vector control RNAi samples to minimize the chance of failing in vector control experiment, which would impact data analysis for all RNAi conditions in the same batch. One vector control sample was prepared side-by-side with RNAi conditions, while the other one was independently prepared in the same day, using separate bacteria culture and bleaching *C. elegans* from a distinct parental animal batch. The latter case aligned with criteria for biological replication (fresh material) except that the experiment was conducted on the same day, thus we refer to them as 'same-day replicates'. We noted that the expression variability between different-day and same-day replicates was similar, therefore, all six vector control samples were treated as biological replicates in differential expression analysis to enhance the statistical power (further details are provided in the following sections).

**Next generation sequencing**

Most WPS sequencing libraries were sequenced on the BGISEQ-500 next-generation sequencer platform with 100-bp paired-end reads. A subset of the libraries was sequenced using an Illumina NextSeq 500 sequencer with a NextSeq 500/550 High Output Kit v2.5 (75 Cycles). For Illumina sequencing, paired-end sequencing was performed with 14 cycles for read 1 and 75 cycles for read 2.

**WPS raw data processing**

The pair-end reads data were either received from BGI directly or produced through standard bcl2fastq procedure with illumina platform. Reads were processed by an in-house dolphinNext pipeline[12] to generate a gene-by-sample read count matrix. The pipeline includes the following steps: (1) raw reads were demultiplexed by a homemade python script that extracts the barcode information from read 1 and combines that with read 2. (2) The processed reads were passed to Trimmomatic (v0.32)[13], to remove polyA and adaptor sequences. (3) next, reads were aligned to the *C. elegans* genome (WormBase WS279) by STAR[14] (parameter: *--runThreadN 4 --alignIntronMax 25000 --outFilterIntronMotifs RemoveNoncanonicalUnannotated*). (4) Finally, the output bam file was processed by ESAT[15] to obtain the read counts of genes. In ESAT, we used an extension window of 1000 bp and the 'proper' method of multiple mappings. Unique Molecular Identifier (UMI) features were not used by setting *umiMin = 1*. Read counts, rather than UMI counts, were used as the gene expression quantity. We did not observe significant PCR duplicates during the development of our method (i.e., read counts highly correlate with UMI counts, data not shown), which is consistent with the low number of PCR cycles for sequencing library construction (**Supplementary Protocol**). Therefore, we directly used the read counts regardless of the presence of UMI in our sequencing library. The dolphinNext pipeline also includes a few quality control (QC) procedures for sequencing library and alignment quality and is interactive through the online portal[12]. The pipeline processes each WPS sequence library individually and produces a read count table for the sequencing library. The pipeline can be downloaded at https://github.com/XuhangLi/WPS.

Reads count tables were used as the input for all downstream analyses. Reads from ribosomal RNA (i.e., mapped to ribosomal genes) were discarded. The sequencing library depth was measured with the sum of read counts of each sample after ribosomal gene removal. Samples with depth lower than 1 million were removed.

**WPS RNAi identity QC and dsRNA decontamination**

We discovered that the reads from dsRNA in RNA interference (mostly in anti-sense strand) could be used to determine the identity of the RNAi clone used. However, these reads might potentially confound the quantification of the RNAi target gene expression since some map to the sense strand at the 3' end of the gene. In rare cases, they can also influence the quantification of other genes when their transcripts extend to regions containing dsRNA reads. Therefore, we developed a python script to both quantify the dsRNA (anti-sense RNA) signals and the expression levels of the dsRNA-influenced genes. This is feasible because dsRNA signals are confined to the coding region of the target gene, while the mRNA signals predominantly reside at the 3'-UTR of the transcript. We achieved it by identifying genomic regions covered by dsRNA signals and re-quantifying genes that were influenced by only counting the reads in the clean regions.

We performed the dsRNA analysis on a library-by-library basis, ensuring that any re-quantification of gene expression was uniformly applied to all samples within a sequencing library. To identify possibly dsRNA-influenced genes, we searched for genes whose transcripts overlapped with the exons of any RNAi target gene in the sequencing library. This gave a set of genes to be corrected for potential dsRNA contamination. Next, we identified the genomic regions contaminated by thresholding the reads mapped to the complementary strand of the mRNA for each RNAi-targeted gene. Finally, the read counts of all potentially contaminated genes were recounted using the clean (not contaminated) regions only.

dsRNA signals were quantified by counting reads mapping to the complementary strand of the mRNA(s) for each RNAi-targeted gene. This dsRNA quantification procedure was applied to all metabolic genes to identify the potential cross-contamination and sample swaps. When applicable, the procedure was applied to a control bam file that was made from an RNA-seq library of animals treated with only vector control, establishing the background level of reads mapping to the complimentary strand for each gene and was used to calculate the enrichment of dsRNA (anti-sense RNA) signal in the RNAi identity QC. The control sequencing library used in our study was the developmental stage sequencing library (see the corresponding section below for details).

Since the dsRNA-influenced genes were quantified solely by reads mapped to the clean region, it may significantly reduce the total reads (depth) for a gene, potentially resulting in a loss of power. Therefore, we applied such dsRNA de-contamination only to genes whose loss of depth was less than 50% (recounted read counts in vector controls were greater than or equal to 50% of the original read counts). Genes that were not corrected were noted, and additional scrutiny was applied when evaluating their RNAi efficiency. The dsRNA (anti-sense RNA) analysis is available in WPS data analysis pipeline (https://github.com/XuhangLi/WPS).

**WPS RNAi efficiency QC**

We performed QC of RNAi efficiency based on two complementary criteria: the reduction of reads for the targeted gene and/or the detection of target anti-sense RNA. A reduction in reads for the targeted gene may not always be observed even if the RNAi is successful because the gene is lowly expressed or if the expression quantification is influenced by dsRNA and cannot be decontaminated (see above). Therefore, we considered an RNAi-condition to pass QC when there was either a two-fold decrease in reads of the targeted gene and/or a greater than 10-fold increase in anti-sense RNA signals corresponding to the targeted gene. To simplify this quantification, we calculated the fold change simply by dividing the TPM of the targeted gene in the RNAi condition by that in the vector control condition within the same batch. Similarly for anti-sense RNA signals, we divided the anti-sense RNA count-per-million (CPM) in a RNAi condition by the background anti-sense RNA CPM based on a vector-control-only sequencing library that was described in the previous RNAi identity QC section.

We also used the anti-sense RNA signal to identify potential cross-contaminations (i.e., one condition contains anti-sense RNA mapping to two genes). Such cross-contaminated samples were rare and were either labeled as 'MULTIPLE' in the sample metadata and included in the dataset or removed. Together, the QC pipeline outputs a list of failed-QC conditions and evaluation figures (such as the heatmap of anti-sense RNA) for manual interpretation. All QC results were carefully inspected to ensure the quality of the dataset.

For any RNAi conditions that did not pass RNAi QC, we performed Sanger sequencing of the RNAi clone. We found these fail-QC RNAi carried plasmids that (1) contain an insert lacking at least 100 consecutive base pairs targeting to a *C. elegans* gene (‘SHORT); or (2) contain an insert that do not target to any *C. elegans* genomic region (‘VECTORLIKE’); or (3) contain a recombined vector that lacks the T7 promoter, therefore deficient in expressing dsRNA (‘RCBVECTOR’); (4) contain an insert that targets to multiple *C. elegans* gene (‘MULTIPLE’); (5) undefined RNAi identity because the Sanger sequencing did not return a signal (‘NOSIGNAL’), or (6) contain the RNAi insert that targets to another *C. elegans* gene. The last were relabeled in the metadata table and included in the final dataset. The erroneous RNAi such as short inserts were relabeled with specific prefix (e.g. ‘SHORT_’) in the sample name and was used in the analysis when applicable (e.g. forming the set of non-targeting perturbation (NTP)).

As a showcase of the frequency of these fail-QC perturbations, we found among the 3,784 samples generated in the metabolic WPS experiment, 76 (2.0%) were removed due to low depth (< 1 million) or bad quality (see below), 89 ‘SHORT’ (2.4%), 71 ‘VECTORLIKE’ (1.9%), 38 ‘RCBVECTOR’ (1.0%), 33 ‘MULTIPLE’ (0.9%), 20 ‘NOSIGNAL’ (0.5%) and 254 (6.7%) swapped to targeting another *C. elegans* gene. Together, the on-target pass-QC rate for a large-scale WPS is expected to be ~85% (3,203/3,784) including vector control samples.

**WPS sample quality QC via exploratory data analysis (EDA)**

To identify the 'outlier' samples, we performed library-level EDA based on a serial manual inspection of plots based on Principal Component Analysis (PCA), Euclidean distance and Pearson correlation, which is a common practice for RNA-seq analysis (http://bioconductor.org/packages/devel/bioc/vignettes/DESeq2/inst/doc/DESeq2.html). We consider a sample to be problematic ('bad sample') if it displayed high distance to other replicates (usually > 50), existed as a clear outlier in PCA plots, and/or showed poor sample-sample correlation (r-squared < 0.95) or had large set of outliers in the inter-replicate gene expression scatter plot. A bad sample usually satisfies most or all of these criteria. We automated the generation of these QC plots but did not automate the identification of bad sample. We reasoned that samples may go wrong in different ways such that the thresholds for one study/experiment may not be applied to another. For instance, in the metabolic WPS data, we noticed that a sample could be an outlier in PCA plot and show significant distance with other replicates, however, displayed good correlation (i.e., $r^2 > 0.98$) with other replicates. Further investigation found that this is due to the difference in their sequencing depths (i.e., the one is less than 2 million). Therefore, we did not consider such samples as bad samples. Since being interactive is the nature of EDA, this part of QC was designed to require manual inspection of the data by the researcher. This is not a speed limiting step of the data processing as inspecting the plots of one sequencing library usually only takes a few minutes. Of note, bad samples are rare in WPS routine, for instance, only 48 (1.3%) samples were identified in the metabolic WPS dataset.

#### Control-dependent differential expression (DE) analysis

Typically, a sequencing library includes 15-16 RNAi conditions in triplicate and six vector control samples. As mentioned above, these six vector control samples were collected over three different-day replicates, each compromising two independently cultured, same-day replicates. We initially analyzed the gene expression variance level within the two same-day replicates and found it was similar to that among the different-day replicates (**Supplementary Protocol**). In addition, we noted that DE analysis solely based on different-day replicates often produced slightly more DEGs compared with using all six samples (data not shown). This may be because it is less prone to underestimating variations with a greater sample size. Together, we reasoned that since using six replicates practically generates more conservative results, and theoretically can be more powerful because of increased sample size, we decided to use the six control samples as six biological replicates in our WPS DE analysis. We acknowledged that the two kinds of replicates may behave differently in the hand of another researcher, so we advise WPS users to carefully evaluate before deciding on using only different-day replicates versus all the six (see **Supplementary Protocol**).

The control-dependent DE analysis was performed using DESeq2 (v1.26.0[16]). Given that sequencing library construction can introduce batch effects, we conducted DE analysis on a per-library-basis using roughly 50 samples in each run of DESeq2. To mitigate potential batch effects between replicates, replicate batch information (i.e., rep1, rep2, and rep3) was incorporated into the DE model (*design ~ replicate_batch_label + RNAi_condition_label*). Genes with fewer than 10 read counts across all samples in a sequencing library were excluded from the DE analysis. We disabled independent filtering (*independentFiltering = F*) and instead employed a custom filter (see below) for consistency across sequencing libraries. We produced two log-fold-change estimates, including *log2FoldChange* estimates from DESeq2[16] (referred to as raw fold change) and the shrinkage estimates from *apeglm* (*apeMethod='nbinomC'*) (referred to as shrunk fold change), which were compared and utilized as descried in the section *WPS analysis parameter selection*.

Together, this control-dependent DESeq2 analysis follows standard procedures of DESeq2 and is also referred to as DESeq2 approach (as compared with EmpirDE approach) or conventional DE analysis in this paper. The outputs here form a foundation for further test statistic modeling in the EmpirDE analysis.

**Control-independent differential expression analysis**

The idea of control-independent DE analysis is to perform DE analysis by comparing an RNAi condition against all the other samples within the same sequencing library. Given that DE is typically sparse and condition-specific in large scale screening, we expect most if not all genes will be affected, and hence exhibit differential expression, in only a limited number of conditions within a sequencing library (i.e., DE call percentage < 30%, meaning the frequency of DE call for any gene is less than 5 out of 16 conditions in a library). Consequently, most conditions in a sequencing library can serve as a null population for DE analysis that does not rely on control labels.

However, if a gene is truly differentially expressed in multiple conditions within the same sequencing library, the power to detect DEGs will be reduced when one condition is compared directly with all others. A more effective approach is to compare an RNAi condition only with a true null population, in which the gene of interest is not differentially expressed. This approach, however, poses a challenge of identifying the main (null) population based on gene expression data. We used AdaTiss [6] for robust fitting of the gene expression across all samples and to exclude samples in which the gene's expression was extreme with respect to this fit. With size-normalized and batch-corrected expression levels (corrected using the *removeBatchEffect* function in *limma* package), we applied AdaTiss to fit the mean and variance for each gene in each sequencing library, one at a time (example command: *out = AdaReg(model.matrix(~1,data = as.data.frame(y)), y), where y represents the expression level vector*). A fit was deemed successful if $pi_0 \geq 0.7$ (at least 70% of samples were in the main population), and was then used to calculate z-scores for each condition (*z = (y-out$beta.rob.fit)/sqrt(out$var.sig.gp.fit)*). Overall, the rate of successful AdaTiss fitting is usually around 95%. In the case of unsuccessful fitting, we used simple statistics as a surrogate. For genes lowly expressed (median normalized read count ≤ 10), we used the mean and standard deviation to calculate z-score. For those highly expressed ones (median count greater than 10), we used the median and mad (median-absolute-deviation). The rationale for using mean/sd for lowly expressed genes and median/mad for highly expressed ones is to mitigate the high variance for lowly expressed genes (thus, mean/sd provides a more conservative estimation of the main population) while maximizing the power for highly expressed genes.

To define the null population based on the fitting, we used a z-score cutoff of 2.5 (equivalent to a *P* value of approximately 0.01). For each gene, any condition (including vector controls but excluding the specific RNAi under analysis) with a median z-score (across three replicates) below 2.5 or above -2.5 was included in the null population. Conditions not meeting this criterion were categorized as 'outlier' population. In the uncommon event where over 50% of conditions in a sequencing library were identified as outliers, making it likely that many RNAi affected the gene, we conservatively designated all conditions within that library as the null population to reduce the risk of false positives in such scenarios.

To streamline DE analysis with DESeq2, i.e. to build a DE model using a single gene expression matrix, we replaced the outlier expression values with imputed values derived from inliers (the null population). This strategy was adopted to circumvent the need to run DESeq2 separately for each gene because of different null populations, which would be computationally impractical without modifying the DESeq2 package. A bootstrap strategy was used for imputation. For each expression value in outlier conditions, we randomly selected a corresponding expression value from inlier conditions in the same biological replicate. To do so, we use the DESeq2 normalized counts for inlier samples and multiply the sampled value by the outlier sample size factor, rounded to an integer, to get an imputed count value. This procedure resulted in a new read count matrix, wherein the values for outlier conditions of each gene were replaced with these imputed counts, based on inliers identified through robust-fitting z-scores.

Like the control-dependent DE, genes with fewer than 10 read counts across all samples in a sequencing library were excluded from the analysis of that library. To manage potential single outlier samples within the null population, we enabled the outlier replacement function in DESeq2 by setting *minReplicatesForReplace=7*. Other DE parameters were identical to those used in vector-dependent DE analysis. DE results were derived by contrasting the targeted RNAi condition against the defined null population.

#### Combining control dependent and independent DE analysis results in the EmpirDE framework

The EmpirDE analysis integrates results from both control-dependent and independent DE analyses to resolve problems caused by control-outlier genes (**Fig. 3d**). This is necessary because control-independent DE analysis can be unreliable when the null population is inaccurately estimated. Therefore, we combined the control-dependent and independent DE results to optimize power and error rates in EmpirDE. The approach involves applying control-independent DE analysis solely for control-outlier genes.

We developed a single parameter, the outlier threshold (*P_out*), a *P* value cutoff, to determine whether a gene should be regarded as a control-outlier gene. The control-outlier genes were identified based on two criteria: (1) within a sequencing library, this gene was unidirectionally (i.e., either all increased or decreased) differentially expressed in at least 50% of RNAi conditions in control-dependent DE analysis with a *P* value below the threshold of *P_out*; and (2) concurrently, at least 75% of RNAi conditions were coherently differentially expressed under a relaxed threshold of *P_out * 10*. By managing the 50% and 75% quantiles, this approach pinpointed genes where the overall RNAi conditions shifted up or down in gene expression compared to the vector control. By default, EmpirDE used a *P_out* of 0.005, whose determination is described in the following section *EmpirDE parameter selection*.

In each sequencing library, we applied control-independent DE results to all identified control-outlier genes. There were a few additional considerations. First, if any gene was found to be differentially expressed with substantially greater statistical significance in control-independent DE analysis – defined by *P* values at least 100 times lower and a higher fold-change – control-independent DE results were used to enhance the power of DEG discovery. Second, to maintain consistent empirical null modeling (see details below), genes marked as control-outlier genes in more than 25% of libraries (e.g., for metabolic WPS, this is 72 libraries * 0.25 = 18) had control-independent DE results applied across all libraries. This was regardless of whether they were identified as outlier genes in each individual library.

**Empirical null modeling of DE test statistic**

The empirical null was modeled individually for each gene by combining all conditions in a WPS experiment (dataset). In a standard WPS application, at least 96 conditions are experimented, providing a substantial sample size for building the empirical null.

We used the fitting function of the *locfdr* package in R to model the empirical null. For each gene, its Wald statistics generated by DESeq2 across all experimental conditions (> 100) were used as the input for the *locfdr* function. The command used was: *locfdr(target_gene_wald_statistics, bre = brk, plot = 0, type = 0)*, where *brk = length(target_gene_wald_statistics) %/% 8*. This break size (*bre*) formula was empirically determined based on what gave the best fit in manual inspections. Extreme outliers in the Wald statistic (defined as greater than the 99% quantile plus 3 MAD (Median Absolute Deviation) or less than the 1% quantile minus 3 MAD) were excluded from the fitting, as the presence of such outliers could cause the program to fail. Occasionally, *locfdr* would exit with an error due to issues in fitting the distribution. In these cases, we incrementally increased the break size (bre = brk + 1, 2, 3, ...) until a successful fit was achieved. In rare situations where fitting could not be completed after 100 increments, we used the median and MAD of the Wald statistic distribution to estimate the empirical null. Upon determining the null's parameter estimates, we rescaled the Wald statistic to compute a corrected Wald statistic and subsequently calculated the new *empirical P* values (**Fig. 3d**).

To compute the empirical FDR, we applied a bi-directional multiple testing correction to conservatively control the false discovery rate. This strategy was also benchmarked through simulations (see below for details). To increase power and exclude very lowly expressed genes (akin to independent filtering in DESeq2), we first filtered the genes with median normalized counts in both vector control and RNAi samples of 30 or less (individually for each DE comparison). These filtered genes were assigned with an adjusted *P* value of *NA*. To adjust for multiple testing, a row-wise adjusted *P* value was calculated using the Benjamini-Hochberg (BH) method across all conditions for a given gene. Simultaneously, a column-wise adjusted *P* value was calculated using BH method across all pass-filter genes for a given condition. We defined the empirical FDR as the maximum of the row-wise and column-wise adjusted *P* values. This worst-case FDR approach ensures that the rate of false DE calls among all genes for a given RNAi condition, and the rate of false calls among all conditions for a given gene, are both below the desired threshold.

**WPS data simulation**

To mimic the real metabolic WPS dataset collected from 72 WPS sequencing libraries across 12 RNAi plates, we used scDesign3[17] to simulate each batch, i.e., each sequencing library, individually. A typical sequencing library contains 16 RNAi conditions in triplicates and 6 vector control samples. In the simulation, we first removed lowly expressed genes with a maximum read count of 10 or less. The read count matrix was then used to estimate simulation parameters via *fit_marginal* function in scDesign3. We incorporated the RNAi condition as the sole covariate in fitting mu (*mu_formula* = *'condition'*) and bypassed the marginal distribution fitting of standard deviation (*sigma_formula* = *'1'*). The canonical negative binomial model was used throughout (*family_use* = *'nb'*). Marginal distribution estimates were subsequentially input into *fit_copula (copula* = *'gaussian'*) to determine gene correlation parameters. These together established the simulation parameters for a sequencing library.

To simulate differential expression, we defined the ground truth for DEGs using a total of 117,782 DEGs identified in real data by the default WPS data analysis method

(FDR < 0.1, FC > 1.5). This ground truth table decides which genes in which RNAi conditions should be simulated as DEGs and their desired fold changes. Next, we reconstructed the mean estimate matrix from the parameter estimation step to reflect the DEGs to be simulated. To achieve this, we first calculated the average fitted mean for each gene using the mean matrix to define its *reference expression level*. For genes designated as differentially expressed, we defined their new means in the reconstructed mean matrix as their reference expression levels multiplied by the desired fold changes from the ground truth table. For other genes, their new means were simply defined as the reference expression level, simulating no differential expression. Finally, a simulated sequencing library was generated using *simu_new* function, employing the reconstructed mean matrix and other estimated parameters as input.

In simulations incorporating $\Delta\mu$, we added random noises ($\Delta\mu$) to the reconstructed mean matrix before generating simulated data. We empirically determined the individual level of random noise for each gene based on comparisons between real data and standard-NB simulated data (the simulation data generated without adding $\Delta\mu$, as stated above). Specifically, we first identified inflated genes that required delta μ addition to align with real data (**Extended Data Fig. 2c**). These are genes whose log2(FC) variation across all conditions (also see below) was greater in real data than in standard-NB model simulations (**Extended Data Fig. 2c,** $\sigma_{real} > \sigma_{NB}$, referred to as inflated genes). A random delta μ was then added to these inflated genes using a heuristic formula that best captured gene-specific mean fluctuation (**Extended Data Fig. 2c, Equation 1-2**). Notably, this random delta μ was added to the means of inflated genes, irrespective of their differential expression status.

$$log2(FC_{\Delta\mu}) = N(0, (0.8 \times \sqrt{{\sigma_{real}}^2 - {\sigma_{NB}}^2})^2) \quad \textbf{Equation 1}$$

$$\Delta\mu = \mu_0 \times (FC_{\Delta\mu} - 1), \mu_0 \ is\ the\ reference\ expression\ level \quad \textbf{Equation 2}$$

**Benchmarking EmpirDE using simulation data**

The DE analysis on simulated data follows the same procedures as those used for real data analysis stated above. There were a few modifications because of the differences between real and simulated data. Firstly, control-independent DE analysis was omitted because control-outlier effects were not simulated, given that it is irrelevant to the analysis of empirical null modeling. Therefore, for empirical null fitting, we directly used DE results from control-dependent DE analysis. Secondly, we excluded DE analysis for two sequencing libraries (met2_lib5 and met3_lib4) because they either lacked three replicates pooled in the same library (met2_lib5) or were without vector controls (met3_lib4) (for details on these two libraries, see [REWIRING]). This reduced the total number of DEGs used for benchmarking analysis to 117,096. Thirdly, we implemented a simplified DE model using only the RNAi condition as covariate, as batch effects in replicates were not simulated. Other procedures remained consistent for both DESeq2 and EmpirDE analysis.

Using DE analysis outputs, we analyzed the distribution of log2(FC) in both simulated and real metabolic WPS data to derive parameters ($\sigma_{real}$ and $\sigma_{NB}$) for delta μ modeling. The log2(FC) distribution for each gene was fitted following the same method as that for fitting the empirical null with the Wald statistic, to produce the corresponding $\sigma$. To increase the robustness of this parameter estimation, we performed standard NB simulations of WPS data 10 times and averaged the $\sigma$ generated for each gene to determine its parameter $\sigma_{NB}$.

We benchmarked EmpirDE performance with the pool of 117,096 ground truth DEGs across the entire dataset. For each FDR threshold (ranging from 1e-30 to 1), we calculated the actual False Discovery Proportion (FDP, FP/(FP+TP)) and power (TP/(TP+FN)) based on DEGs solely defined by the FDR threshold. The metabolic WPS data were independently simulated 10 times to evaluate variability in FDP and power. To compare multiple testing adjustment strategies, we applied BH adjustments post-filtering of lowly expressed genes, as stated above. The lowly expressed genes were those with median normalized counts below 30 in both control and the RNAi condition (approximately equivalent to 5 in TPM). ‘Column-wise’ adjustment refers to adjusting multiple tests for genes within a single condition, using *P* values of all pass-filter genes in that condition. ‘Row-wise’ adjustment was for multiple tests across different conditions for a specific gene, using that gene's *P* values across 1,078 conditions in this simulation. The worst-case adjustment took the higher adjusted *P* values from these two approaches.

#### Benchmarking EmpirDE using non-targeting perturbations (NTPs)

We identified 71 RNAi conditions, including the four spike-in controls, as non-targeting perturbations (NTP) based on the RNAi identity QC and Sanger sequencing. These RNAi clones failed RNAi identity QC, lacking substantial dsRNA detection and targeted gene knockdown, and were found to be either 'SHORT', 'VECTORLIKE', or 'RCBVECTOR' in Sanger sequencing. Thus, they should not effectively target any *C. elegans* genes, and can serve as independent negative controls (**Supplementary Table 2**). We used the number of DEG calls in these conditions to evaluate false discoveries. Given an FC and FDR threshold, we used the 90% quantile of the number of DEGs in these 71 conditions as an estimate of false positive calls. Choosing the 90% quantile, rather than the median, mean or max, aims to be conservative ('overestimating' false positives) while also allowing for a few outlier conditions that might not act as true negative controls.

#### Benchmarking EmpirDE using reproducibility

A total of 36 RNAi conditions were independently experimented 2-3 times (each with three replicates), serving to assess the reproducibility, which in turn provided a proxy of false positives. The strictly unreproducible DE calls were considered as empirical false positives. These were defined as DEGs in one experiment (FDR<0.2, FC>1.5 for **Extended Data Fig. 3e** and FDR < 0.1, FC > 1.5 elsewhere; using a slightly relaxed FDR threshold in **Extended Data Fig. 3e** was to include more DEG calls for better evaluating parameters) but the gene showed no significant change in expression in another (FC < 1.1) or exhibited a reverse expression change (e.g., one increased while the other decreased). This approach offers a qualitative assessment of DEG reproducibility, thus serving as a proxy for false discoveries in DE analysis. The unreproducible DEG rate was calculated as the number of strictly unreproducible DEG calls divided by the total DEG number of that experiment. The average rate of the two repeats is shown in **Fig. 4f**.

Of note, two of the 36 conditions had more than two (i.e., three) independent repeats. We conducted all pairwise comparisons among the three (resulting in three combinations). Therefore, a total of 40 pairs were used and displayed.

**EmpirDE parameter selection**

We systematically evaluated parameters in the EmpirDE framework, using NTPs and independent repeats. These included the *P_out* threshold for control-outlier gene selection, two FC estimates (log2(FC) from DESeq2 and the shrunk log2(FC) from *apeglm*), and thresholds to define DEGs (FC and FDR thresholds).

Evaluation using the 71 NTP conditions showed that false positives were insensitive to the type of FC estimates when the FDR cutoff was adequate (e.g., FDR < 0.1, data not shown). A similar observation was made with the 36 repeats. Given that FC estimates from DESeq2 yielded slightly more DEG calls (~5%) and are simpler, we opted for this FC in our standard EmpirDE analysis. Accordingly, this FC was also used when reporting the result of DESeq2 analysis (**Fig. 3a**).

Regarding *P_out*, we evaluated thresholds ranging from 0 to 1, and found that the values between 0.001-0.01 were optimal when evaluated with NTP conditions (**Extended Data Fig. 3d**). A similar optimal range (0.0025-0.0075) was observed with the 36 repeated conditions (**Extended Data Fig. 3e**). Thus, we selected 0.005 as the *P_out* parameter, which is ten-fold lower than the common statistical threshold of 0.05 but is reasonable because this threshold is on *P* values without multiple testing adjustments.

Regarding the thresholds for defining DEGs, we reasoned that an FDR of 0.1, expecting 10% false discoveries, is ideal to balance power and error, given that benchmark analyses demonstrate the FDR in EmpirDE approach is statistically rigorous (**Fig. 4**). We then selected a fold-change threshold (1.5) that controlled false positives in NTPs below five under this FDR threshold (**Fig. 4d**). This fold-change threshold can further eliminate false discoveries and uninteresting DEGs that only changed subtlety.

Together, we settled on a parameter combination of $P$_out = 0.005, empirical FDR < 0.1 and DESeq2 fold change estimate greater than 1.5 to define the final DEGs. This parameter set demonstrated stringent FDR control and established a responsiveness cutoff of 5 DEGs.

**Alterations in experimental setup of NHR WPS**

The NHR WPS dataset was generated during the early stages of this project, and due to historical reasons, has a different sequencing library design. Consequently, data processing was slightly altered to accommodate.

The primary distinction lies in the arrangement of the sequencing library. In the NHR experiment, samples from the same biological replicate were pooled together in a sequencing library, comprising 47 RNAi conditions and 2 vector controls. Accordingly, the three replicates of a RNAi condition were sequenced in three separate sequencing libraries, which is different from the metabolic WPS. Most NHR data were collected from an experiment using a single 96-well RNAi plate, producing two sequencing libraries that contained vector control samples derived from the same total RNA extraction. Consequently, the vector controls in the two sequencing libraries were technical replicates and should not be included simultaneously in DE analysis. The majority of NHR data were from 6 sequencing libraries (3 replicates x 2 libraries each), and there were two supplementary libraries created to incorporate extra NHR conditions and to redo the experiments for some conditions that failed quality control. This led to a total of 104 unique NHR RNAi conditions with 100 out of the 104 conditions assessed in triplicate or more (up to five biological replicates). The remaining four were in duplicate.

Lastly, the construction of NHR sequencing library followed an earlier version of WPS library construction protocol. A notable difference is the use of SuperScript™ II instead of SuperScript™ III that is used in standard WPS (**Supplementary Protocol**). However, we did not notice obvious differences in the detection sensitivity of genes driven by this change of reverse transcriptase (data not shown).

#### NHR WPS QC

Quality control (QC) for NHR WPS dataset was conducted in a manner similar to the metabolic sequencing library. However, it is important to note that the NHR sequencing library was collected in the initial phase of the project, a period when many methodology optimizations were still ongoing, the pass-QC rate for the NHR sequencing library is somewhat lower than that observed for the metabolic sequencing library. Out of 392 samples, a total of 353 (~90%) passed quality control. The fail-QC samples include 19 bad quality samples (5%), a frequency notably higher than metabolic sequencing library (2%), and 20 failed RNAi identity QC due to a pipetting error that caused cross-contamination.

#### NHR WPS DE analysis

The DE analysis for the NHR sequencing library posed unique challenges due to its distinct library arrangement, necessitating a different DE strategy rather than the within-library analysis. We divided all samples (across 8 sequencing libraries) into four DE groups to maximize sample pooling, which aids in better estimating dispersion, and to prevent the co-occurrence of technical replicates of vector controls. The four DE groups were defined as follows: Group 1 included most conditions from the first half of the 96-well plate, along with additional conditions from the supplementary library #1. Group 2 comprised the remaining conditions from the first half of the 96-well plate, plus their extra replicates sequenced in supplementary library #2. Group 3 contained most conditions from the second half of the 96-well plate, plus their extra replicates from supplementary library #1. Group 4 involved the remaining conditions from the second half of the 96-well plate, plus their extra replicates from supplementary library #2. This grouping strategy was complex and a compromise, reflecting the less refined experimental design at the early stages of the project. This issue was unique to the NHR dataset.

The standard EmpirDE analysis was applied to the NHR dataset, with control-dependent and -independent DE performed using samples from the four DE groups separately. The parameter choices remained consistent with those used in the rest of the project. Additional modifications specific to the NHR dataset are as follows:

1. We noted that the NHR sequencing library exhibited a significantly lower incidence of control-outlier genes compared to the metabolic WPS (~10-fold less, data not shown). We suspect a relevance to the preparation of bacterial diet: for NHR WPS, control bacteria were cultured in the same 96-well plate with the RNAi bacteria. However, for metabolic WPS, control bacteria were cultured in a second 96-well plate because of the inclusion of more RNAi conditions in each experiment (usually ~120 conditions, **Supplementary Protocol**). Based on this observation, we proposed an optimized experimental design in the **Supplementary Protocol** for future users.
2. A comparison of empirical nulls between the metabolic and NHR WPS revealed distinct, albeit moderately correlated, standard deviations (data not shown), suggesting each experiment exhibits unique noises to address. Therefore, the gene-specific random fluctuations can be influenced by experimental batches, and the empirical null needs to be reconstructed in each individual WPS study for its best outcome.

**Transcriptional profiling of animals in different developmental stages**

We profiled the transcriptome of animals at different developmental stages, ranging from L2 larvae to adult, using the WPS sequencing library construction method. This includes 51 samples from animals at 17, 25, 35, 40, 45, 50, 55, 60, 61, 62, 63, 64, 65, 66, 67, 68, 69 hours post L1 plating in biological triplicate. Animals were fed on HT115 bacteria expressing empty vector control (L4440), i.e., the dsRNA expression was induced as in regular WPS experiments. This is to be aligned with the dietary condition of WPS study. The raw data were processed following standard WPS procedures to obtain the read count matrix.

For methodological development purposes, we constructed two technical replicates of this sequencing library using different reverse transcriptases (SuperScript™ II and III). The library generated by SuperScript™ III was used as the non-RNAi control in the RNAi identity analysis of standard WPS (for RNAi identity analysis, see above sections). Analysis of the two sequencing libraries did not find notable differences in detecting gene expression other than batch effects. Therefore, for downstream analysis, we aggregated data from the two replicates by adding up read counts. In five samples (55h_rep1, 68h_rep1, 67h_rep1, 63h_rep3, and 55h_rep2), we noted a slight decorrelation with their biological replicates, a deviation from the correlation levels observed in other samples. This decorrelation can be related to variations in development rate or sample quality in these specific samples. To ensure the integrity of data interpretation, these five samples were excluded from downstream analysis. Sample-to-sample Pearson correlation coefficients were calculated using variance stabilized read counts obtained through the *vst* function in DESeq2 package[16]. These coefficients were then visualized using the *pheatmap* from pheatmap package in R.

The raw and processed sequencing data are available at Gene Expression Omnibus (GEO) session GSE255865. Sequencing libraries constructed with SuperScript™ II and III were made available separately for reference.

**Subsampling analysis for optimal sequencing depth**

To generate a reference profile for subsampling analysis, we aggregated WPS data (i.e., adding up read counts) from one replicate of the developmental stage samples collected at 62, 63, 64 and 65 hours. This produced a transcriptome profile with a sequencing depth of 53 million reads. We used R package *subSeq*[18] to subsample this reference data 10 times at 10 different depths (50 M, 40 M, 30 M, 20 M,10 M, 8 M, 6 M, 4 M, 2 M and 1 M reads). The coefficient of variation (CV) for each gene at each depth was then calculated. Genes with a CV < 0.15 at a specific depth were classified as quantified at that depth. This CV cutoff represents that the majority of sampling population (~95%, two standard deviations) is within a ±30% interval of the true value, identifying the genes that can be accurately quantified. At each sampling depth, we calculated the fraction of genes quantified at varying TPM threshold (greater than 1, 2, 4, 8 or 16) to determine the sensitivity of gene detection.

We opted for sampling final read counts to better compare effects of different final depths rather than raw read depth, however, a similar result was also observed in the subsampling of fastq files (data not shown).

#### Functional enrichment analysis of NHR GRN

We performed the functional enrichment analysis for NHR perturbations with more than 10 DEGs. DEGs of the RNAi targeted gene were excluded from the analysis. Functional enrichment analysis was performed by *enricher* function from *clusterProfiler* package[19] in R. The universe (*universe* parameter) of the enrichment analysis was defined by all genes (16,245) analyzed in the differential expression analysis. WormCat v2 (Nov. 11, 2021)[20] was used for the gene sets.

#### NHR perturbation-perturbation similarity

Perturbation-perturbation similarity was calculated following the same methodology used in metabolic WPS analysis [REWIRING]. In brief, we used filtered log2(FC), derived by masking log2(FC) of RNAi target genes and non-DEGs to zero, to compute cosine similarity values. These cosine values were used to quantify the perturbation-perturbation similarity. Only responsive perturbations were included in this analysis.

To define the *nhr* pairs based on cosine similarity, we constructed a hierarchical tree by *hclust* function in R using 1-cosine similarity as distance input and 'complete' linkage method. The tree was then cut at its first merge of the leaves, yielding either singleton leaves or clustered pairs (**Fig. 6c**). Using these pairs as clusters, we calculated the silhouette score of each data point by *silhouette* function from *cluster* package. The average silhouette score for all pairs formed a metric measuring the fitness of assigning *nhr* into pairs. To determine the statistical significance, we randomized the NHR GRN 10,000 times using edge swapping approach[21], producing a distribution of average silhouette score with random networks. An empirical *P* value was calculated based on this distribution.

**Comparing NHR perturbation-perturbation similarity with protein sequence similarity**

We analyzed the protein sequence of the longest transcript for 52 *nhr* genes sharing a significant overlap of DEGs with another NHR. These sequences were used as input for Clustal Omega online tool provided by EMBL-EBI[22]. Default parameters were used except for choosing 'yes' for 'DISTANCE MATRIX' and 'No' for 'mBed-like Clustering Guide-tree'. The output distance matrix and percent identity matrix were downloaded and used for the downstream analysis. We also analyzed the DNA biding domain (DBD) sequence similarity of the same 52 *nhr* genes by the same method. The DBD of NHRs were predicted by Conserved Domain Search Service (CD Search) from NCBI with the default settings.

The distance matrix was used to build a hierarchical tree with which the percent identity matrix was visualized in **Fig. 6d**. The percent identity matrix was also used to compare with cosine similarity directly (**Fig. 6e. f**).

**Compare NHR WPS perturbation-perturbation similarity with gene coexpression**

To comprehensively characterize *nhr* gene coexpression, we used a *C. elegans* gene expression compendium that comprises 4,796 samples across 177 datasets[23]. Given that genes can be coexpressed in one dataset but not in another, we first calculated the Pearson Correlation Coefficient (PCC) for gene-gene correlation within each individual dataset. The median PCC in these 177 datasets was then used to quantify the overall strength of coexpression between pairs of *nhr* genes. We compared this median PCC with the corresponding perturbation-perturbation cosine similarity. To quantify the concordance of these two quantities, we focused on the NHR pairs with high cosine similarity (> 0.2) and calculated the median level of their overall strength of coexpression. This observed value was compared to its distribution by random, generated by shuffling the name label of NHR genes in cosine similarity matrix for 10,000 times, to produce an empirical *P* value.

**Supplementary protocol: Worm Perturb-Seq (WPS)**

### Section I: Animal culture and RNA interference

RNA interference library

First, construct WPS RNAi libraries from parent libraries, such as the metabolic[9], TF[10] or ORFome[24] libraries. We recommend constructing a working library for each WPS study using single colonies from the parent library. Specifically, streak the corresponding bacteria onto LB agar plates supplemented with 50 µg/mL ampicillin to allow the formation of single colonies. Pick one colony per RNAi condition, grow the bacteria in 1 mL of LB medium containing 50 µg/mL ampicillin in a 96-deep well plate, and make glycerol stocks in 96-well plates as the WPS RNAi working library.

To reduce the risk of being contaminated, we recommend avoiding placing vector control bacteria directly in the 96-well plates of WPS working library. Instead, make a separate glycerol stock in cryopreservation tube for vector control bacteria. In each WPS experiment, re-introduce the vector control at the step of LB liquid culture alongside RNAi bacteria, either in the same or a different 96-well plate. The details of this process will be discussed in the following section.

RNA interference

RNAi of WPS follows a previously described protocol with slight modifications[11]. Specifically, inoculate bacteria from the WPS working library into a 96-deep well plate with 1 mL of LB medium containing 50 µg/mL ampicillin and shake overnight at 37°C. Dilute 100 uL of each overnight culture by 50-fold using fresh LB medium containing 50 µg/mL ampicillin in a well of a 24-deep well plate and shake for 4 hours at 37°C. After incubation, collect the bacteria by centrifuging the plate at 3,000 g for 20 minutes. Next, resuspend the resulting pellets in 200 µL of M9 solution and transfer to 6-well NGM plates containing 50 µg/mL ampicillin and 2 mM Isopropyl β-d-1-thiogalactopyranoside (IPTG, Fisher Scientific) to induce the expression of RNAi dsRNA. Allow the plates to dry in a hood and then sit on a bench table overnight at room temperature. The RNAi plates are now ready to use for plating worms (see below).

*IMPORTANT: design of an WPS experiment with growth phenotype*

Of note, we generally expect ~10-15% of the RNAi clones to cause a growth phenotype. These conditions can be reassessed by a delayed seeding scheme (see below). However, to always proceed with 96 WPS samples with normal development in each batch of experiment, we recommend two options as follows.

The first option is to start with more than 96 RNAi clones (e.g., 120), by culturing a second plate of RNAi bacteria. This was done in our metabolic WPS screen. However, a substantial batch effect was discovered with this setup, and users should follow our experimental design and computational procedure to eliminate this effect. First, the vector control samples should be cultured in the second RNAi plate, which contains fewer RNAi clones (the ~20 additional clones). This setup ensures the primary population of RNAi conditions is derived from the first plate, forming the null population essential for control independent differential expression analysis to eliminate control outlier genes. Subsequently, users should follow the EmpirDE pipeline for data analysis. If the control bacteria were cultured in the same plate as the main population of RNAi clones, the number of RNAi clones in the second plate would be insufficient to form a null population for control independent differential expression analysis. In such a case, RNAi conditions from the second plate could not be effectively analyzed due to the lack of either same-plate vector control or control-independent null population.

Alternatively, researchers have the option to culture only one 96-well WPS RNAi plate with vector controls in the same plate. In such a case, the batch effect from two plates could be minimized, although it may result in not all WPS samples being sequenced due to the growth phenotype in some samples. As a workaround, a pre-screen can be done to identify and eliminate these phenotypical conditions combined with the delayed seeding scheme, however, at the cost of extra workload.

*C. elegans* culture

Our WPS protocol uses the N2 *C. elegans* strain. To stage the animals properly, follow the animal culturing scheme as below (**Protocol Fig. 1**). For most WPS experiments, plate approximately 200 synchronized L1 animals into each well of the seeded 6-well NGM plate (see **RNAi interference** above) and incubate at 20°C for approximately 63 hours before collection. In the cases where RNAi feeding leads to a developmental delay, feed the synchronized L1 animals with vector control RNAi for 17 hours (i.e., to L2 stage) or 25 hours (i.e., to L3 stage); then transfer the animals to corresponding RNAi plates (**Protocol Fig. 1**). Based on our experience with the metabolic gene RNAi, animals usually develop normally with such delayed RNAi exposure. To perform WPS, only collect animals on which the RNAi treatment does not induce a notable developmental delay.

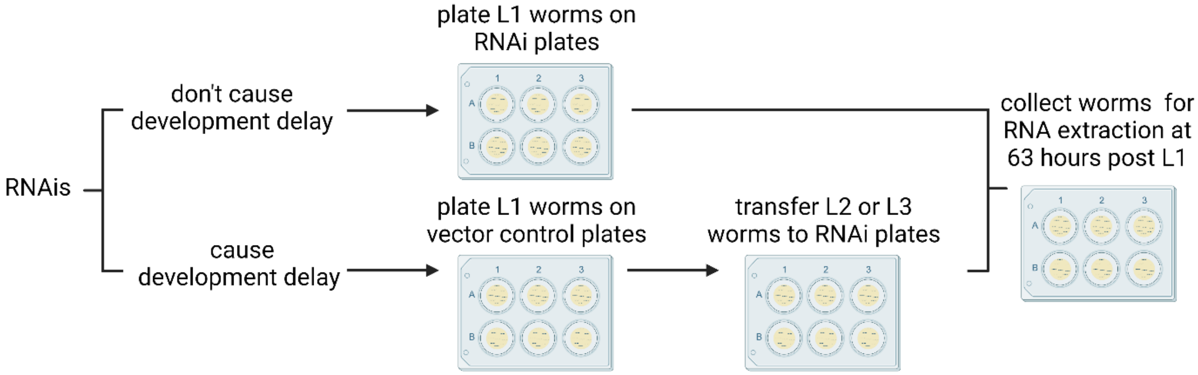


**Protocol Fig. 1**: **Staging of WPS animals**

### Section II: 96-well plate RNA extraction for *C. elegans*

Although the WPS protocol described here extracts RNA from adult stage animals, this RNA extraction protocol can be applied to *C. elegans* from L2 to gravid adult stages. We have not tested this RNA extraction protocol using L1 staged animals. The number of *C. elegans* per sample needed for different stages are listed below.

| Stages | Number of *C. elegans*/sample |
|---|---|
| L2 | >2,500 |
| L3 | >1,000 |
| L4 to young adult | >500 |
| Gravid adult | >200 |

<u>Reagents</u>

1. M9 buffer with 0.01% Triton X-100
2. SDS-DTT solution (200 mM DTT, 0.25% SDS, 20 mM HEPES, pH 8.0, 3% sucrose, stored at −20°C) [1]
3. RNase-free water
4. 100% methanol
5. 100% ethanol
6. 80% ethanol
7. worm lysis buffer [WormBook-Chapter: Reverse genetics. https://www.ncbi.nlm.nih.gov/books/NBK19711/]:
   To make 100 mL worm lysis buffer, mix the following reagents and autoclave:

| **Components and final concentration** | **Stock concentration** | **Qty/100 mL** |
|---|---|---|
| 50 mM KCL | 1 M | 5 mL |
| 10 mM Tris pH 8.3 | 1 M | 1 mL |
| 2.5 mM MgCl2 | 1 M | 0.25 mL |
| 0.01% Gelatin | | 0.01 g |
| $H_2O$ | | 87.85 mL |

The following are added once the autoclaved solution has cooled:

| Components and final concentration | Stock concentration | Qty/100 mL |
|---|---|---|
| 0.45% NP-40 (IGEPAL) | 100% NP-40 (IGEPAL) | 0.45 mL |
| 0.45% Tween-20 | 100% Tween-20 | 0.45 mL |

Store the worm lysis buffer at −20°C and add proteinase K to 1 mg/mL just before use.

8. Monarch® DNA/RNA Protection Reagent [NEB: T2011L]
9. Monarch® RNA Lysis Buffer [NEB: T2012L]
10. Washing buffer 1[3]:

    To prepare 100 mL of Washing Buffer 1, dissolve 11.8 g of Guanidine Thiocyanate (Cat. No. G9277, Sigma) into Rnase-free water to achieve a final volume of 99 mL. Additionally, add 1 mL of 1 M Tris-HCl pH 7.4 (Cat. No. 93313, Sigma) to the mixture. Further pH adjustment is not required. The final concentration of Guanidine Thiocyanate is 1 M now.
11. DNA digestion solution:

    *This solution needs to be freshly prepared before each experiment.*

    For 8 mL:

| |
|---|
| Rnase-free water: 7.1 mL |
| 10 X DNA Digestion Buffer [Fisher: E1010-1-16]: 800 µL |
| Ambion™ Dnase I (Rnase-free) [Thermo Scientific™: AM2222]: 100 µL |

12. 96-Well filter plate (10 µm pore), 960 µL/well [Enzymax: EZ96FTP].
13. 96-Well deep well plate, 2 mL/well [Enzymax: EZ96DWP].
14. 96-Well DNA binding plate, silica membrane [Enzymax: EZ96DBP].
15. AccuBlue® Broad Range RNA Quantitation Kit [Biotium:31073].
16. Greiner Bio-One™ CellStar™ 384-Well, Cell Culture-Treated, Flat-Bottom Microplate [Fisher: 781091].

### Equipment

1. Beckman Coulter Avanti® J-26XP High-Performance Centrifuge with a JS-5.3 Swinging-Bucket Centrifuge Rotor.
2. 1 mL, 300 μL and 10 μL multichannel pipettes.
3. Spark® Multimode Microplate Reader

### Method

(a graphic breakdown is shown in **Protocol Fig. 2**)

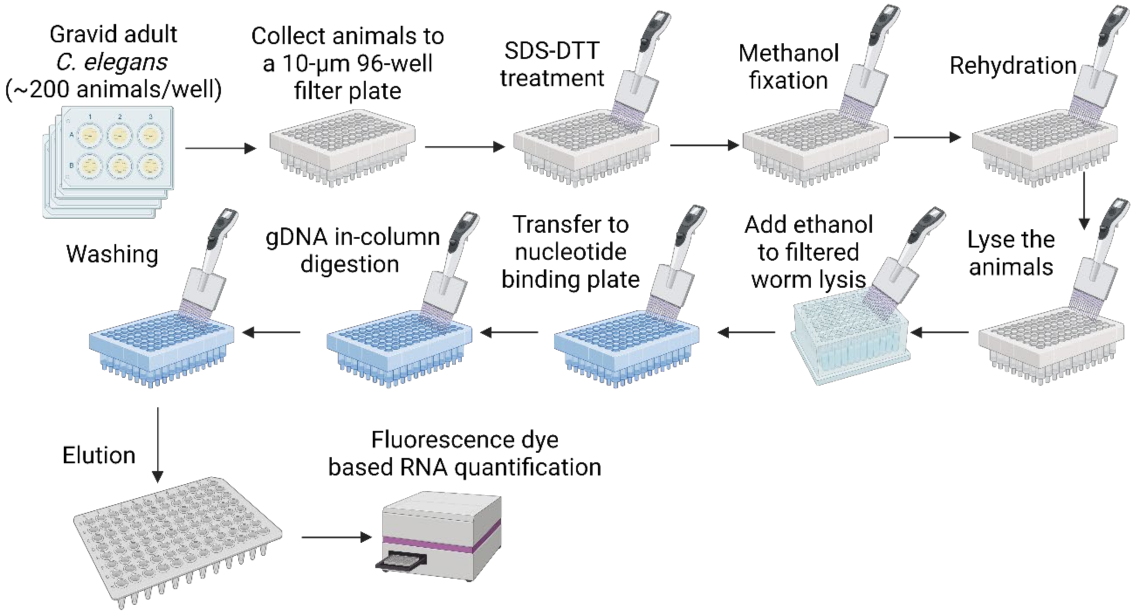


**Protocol Fig. 2. Overview of 96-well plate RNA extraction protocol for *C. elegans***

1. At the time of sample collection (e.g., 63 hours post L1 seeding), wash the animals off the plate with 800 μL of M9 buffer containing 0.01% Triton X-100 and transfer them to a 96-well filter plate. Centrifuge the plate at 2000 g for 2 min and discard the flow-through.
2. Add 100 μL of pre-thawed SDS-DTT solution to each well and incubate at room temperature for 5 min.

3. Add 700 µL of methanol to each well and incubate for 3 min. Centrifuge at 3000 g for 2 min and discard the flow-through. This step is to fix the *C. elegans* tissue and prevent RNA transcriptional changes during RNA extraction.
4. Repeat step 3 two more times.
5. Add 700 µL of $H_2O$ containing 0.01% Triton X-100 to each well and incubate for 3 min. Centrifuge at 3000 g for 1 min and discard the flow-through. Repeat this step two more times.
6. Add 100 µL of pre-thawed worm lysis buffer to each well and incubate at room temperature for 0.5 min.
7. Add 100 µL of Monarch® DNA/RNA Protection Reagent to each well. After adding the reagent, immediately pipette up and down ~20 times. At this point you will see that most tissues are lysed but eggs are still intact when viewed under the microscope. Incubate at room temperature for 5 min. This step is crucial for lysing all *C. elegans* tissue.
8. Add 200 µL of Monarch® RNA Lysis Buffer to each well and incubate at room temperature for 5 min.
9. Put the 96-well filter plate onto a new 96-well deep well plate and centrifuge at 5000 g for 5 min.
10. Add 380 µL of ethanol to the collecting well and transfer liquid to a 96-well DNA binding plate.
11. Centrifuge at 5000 g for 5 min and discard the flow-through.
12. Add 400 µL of Washing buffer 1 to each well. Centrifuge at 5000 g for 5 min and discard the flow-through.
13. Add 80 µL of DNA digestion solution to each well. Seal the top plate and incubate at 37°C for 15 min.
14. Add 400 µL of Washing buffer 1 to each well. Centrifuge at 5000 g for 5 min and discard the flow-through.
15. Add 400 µL of 80% ethanol to each well. Centrifuge at 5000 g for 5 min and discard the flow-through. Repeat this step one more time.
16. Centrifuge the empty plate at 5000 g for 15 min to completely dry the columns.
17. Add 30 µL of Rnase-free water to each well. Place a 96-well PCR plate on top of

the collecting deep well plate and place the DNA binding plate on top of the PCR plate. Centrifuge at 5000 g for 5 min to wash off the RNA. Repeat this step one more time.

18. The elution step usually yields ~30 μL total RNA solution in the PCR plate (the remaining ~30 μL may be retained in the column). Next, measure RNA concentration using the AccuBlue® Broad Range RNA Quantitation Kit in a 384-well plate. Typically, ~30 μL x 200 ng/μL of total RNA can be obtained from each sample.

**Section III: WPS sequencing library construction**

Reagents

1. Nuclease-Free Water
2. TE buffer (Thermo Fisher Scientific: 12090015)
3. Ethanol
4. Qubit reagents: dsDNA HS Assay (Thermo Fisher Scientific: Q32851)
5. dNTP mix (10 mM each) (Thermo Fisher Scientific: R0192)
6. SuperScript™ III Reverse Transcriptase (Thermo Fisher Scientific: 18080044, comes with 0.1 M DTT and First Strand Buffer)
7. RNaseOUT (Thermo Fisher Scientific: 10777019)
8. Second Strand Buffer (Thermo Fisher Scientific: 10812014)
9. DNA Polymerase I (Thermo Fisher Scientific: 18010025)
10. *E. coli* DNA ligase (Thermo Fisher Scientific: 18052019)
11. Ribonuclease H (*E. coli*) (Thermo Fisher Scientific: 18021071)
12. HiScribe™ T7 High Yield RNA Synthesis Kit (NEB: E2040S)
13. Exonuclease I (Thermo Fisher Scientific: EN0581)
14. RNA Fragmentation Reagents (Thermo Fisher Scientific: AM8740)
15. AMPure XP beads (Beckman Coulter: A63880)
16. RNAClean XP beads (Beckman coulter A63987)
17. KAPA HiFi HotStart ReadyMix (Roche: 07958927001)

Equipment

1. Thermocycler
2. Magnetic stand (for 1.5 mL tubes)
3. Qubit® Fluorometer (Thermo Fisher Scientific)
4. Fragment Analyzer (Agilent)

Primers

All our primers maintain the same structure as CEL-Seq2[5], with some differences in the sequences. By using these primers, we can generate sequencing libraries that can be sequenced on both Illumina and BGI platforms.

Barcoding primers

Using primer ‘1S’ (**Table 1**) as an example, the barcoding primer has a T7 promoter sequence (highlighted in red) at the beginning, followed by an Illumina adapter sequence (highlighted in yellow; we replaced the original sequence of CEL-Seq2 with the TruSeq adapter sequence), a 7-nt unique molecular identifier (UMI) sequence (NNVNVNV), the same 6-nt unique barcode as CEL-Seq2 (highlighted in green), and a polyT tail ending with ‘V’.

All our barcoding primers were purchased from Sangon Biotech using polyacrylamide gel electrophoresis (PAGE) purification. The stock solution for these primers is 10 μM in TE buffer, and the working solution is 2.5 μM in TE buffer. We primarily use the first 60 barcoding primers in our experiments as our sequencing library contains less than 60 samples.

**Table 1. The WPS barcoding primers**

| ID | Sequence |
| --- | --- |

| | |
|---|---|
| 1S | GCCGGTAATACGACTCACTATAGGTCCCTACACGACGCTCTTCCGATCTNNVNVNVAGACTCTTTTTTTTTTTTTT<br>TTTTTTTTTTTV |
| 2S | GCCGGTAATACGACTCACTATAGGTCCCTACACGACGCTCTTCCGATCTNNVNVNVAGCTAGTTTTTTTTTTTTTT<br>TTTTTTTTTTTV |
| 3S | GCCGGTAATACGACTCACTATAGGTCCCTACACGACGCTCTTCCGATCTNNVNVNVAGCTCATTTTTTTTTTTTTT<br>TTTTTTTTTTTV |
| 4S | GCCGGTAATACGACTCACTATAGGTCCCTACACGACGCTCTTCCGATCTNNVNVNVAGCTTCTTTTTTTTTTTTTT<br>TTTTTTTTTTTV |
| 5S | GCCGGTAATACGACTCACTATAGGTCCCTACACGACGCTCTTCCGATCTNNVNVNVCATGAGTTTTTTTTTTTTTT<br>TTTTTTTTTTTV |
| 6S | GCCGGTAATACGACTCACTATAGGTCCCTACACGACGCTCTTCCGATCTNNVNVNVCATGCATTTTTTTTTTTTTT<br>TTTTTTTTTTTV |
| 7S | GCCGGTAATACGACTCACTATAGGTCCCTACACGACGCTCTTCCGATCTNNVNVNVCATGTCTTTTTTTTTTTTTT<br>TTTTTTTTTTTV |
| 8S | GCCGGTAATACGACTCACTATAGGTCCCTACACGACGCTCTTCCGATCTNNVNVNVCACTAGTTTTTTTTTTTTTT<br>TTTTTTTTTTTV |
| 9S | GCCGGTAATACGACTCACTATAGGTCCCTACACGACGCTCTTCCGATCTNNVNVNVCAGATCTTTTTTTTTTTTTT<br>TTTTTTTTTTTV |
| 10S | GCCGGTAATACGACTCACTATAGGTCCCTACACGACGCTCTTCCGATCTNNVNVNVTCACAGTTTTTTTTTTTTTT<br>TTTTTTTTTTTV |
| 11S | GCCGGTAATACGACTCACTATAGGTCCCTACACGACGCTCTTCCGATCTNNVNVNVAGGATCTTTTTTTTTTTTTT<br>TTTTTTTTTTTV |
| 12S | GCCGGTAATACGACTCACTATAGGTCCCTACACGACGCTCTTCCGATCTNNVNVNVAGTGCATTTTTTTTTTTTTT<br>TTTTTTTTTTTV |
| 13S | GCCGGTAATACGACTCACTATAGGTCCCTACACGACGCTCTTCCGATCTNNVNVNVAGTGTCTTTTTTTTTTTTTT<br>TTTTTTTTTTTV |
| 14S | GCCGGTAATACGACTCACTATAGGTCCCTACACGACGCTCTTCCGATCTNNVNVNVTCCTAGTTTTTTTTTTTTTT<br>TTTTTTTTTTTV |
| 15S | GCCGGTAATACGACTCACTATAGGTCCCTACACGACGCTCTTCCGATCTNNVNVNVTCTGAGTTTTTTTTTTTTTT<br>TTTTTTTTTTTV |
| 16S | GCCGGTAATACGACTCACTATAGGTCCCTACACGACGCTCTTCCGATCTNNVNVNVTCTGCATTTTTTTTTTTTTT<br>TTTTTTTTTTTV |
| 17S | GCCGGTAATACGACTCACTATAGGTCCCTACACGACGCTCTTCCGATCTNNVNVNVTCGAAGTTTTTTTTTTTTTT<br>TTTTTTTTTTTV |

| | |
|---|---|
| 18S | GCCGGTAATACGACTCACTATAGGTCCCTACACGACGCTCTTCCGATCTNNVNVNVTCGACATTTTTTTTTTTTTT TTTTTTTTTTV |
| 19S | GCCGGTAATACGACTCACTATAGGTCCCTACACGACGCTCTTCCGATCTNNVNVNVTCGATCTTTTTTTTTTTTTT TTTTTTTTTTV |
| 20S | GCCGGTAATACGACTCACTATAGGTCCCTACACGACGCTCTTCCGATCTNNVNVNVGTACAGTTTTTTTTTTTTTT TTTTTTTTTTV |
| 21S | GCCGGTAATACGACTCACTATAGGTCCCTACACGACGCTCTTCCGATCTNNVNVNVGTACCATTTTTTTTTTTTTT TTTTTTTTTTV |
| 22S | GCCGGTAATACGACTCACTATAGGTCCCTACACGACGCTCTTCCGATCTNNVNVNVGTACTCTTTTTTTTTTTTTT TTTTTTTTTTV |
| 23S | GCCGGTAATACGACTCACTATAGGTCCCTACACGACGCTCTTCCGATCTNNVNVNVGTCTAGTTTTTTTTTTTTTT TTTTTTTTTTV |
| 24S | GCCGGTAATACGACTCACTATAGGTCCCTACACGACGCTCTTCCGATCTNNVNVNVGTCTCATTTTTTTTTTTTTT TTTTTTTTTTV |
| 25S | GCCGGTAATACGACTCACTATAGGTCCCTACACGACGCTCTTCCGATCTNNVNVNVGTTGCATTTTTTTTTTTTTT TTTTTTTTTTV |
| 26S | GCCGGTAATACGACTCACTATAGGTCCCTACACGACGCTCTTCCGATCTNNVNVNVGTGACATTTTTTTTTTTTTT TTTTTTTTTTV |
| 27S | GCCGGTAATACGACTCACTATAGGTCCCTACACGACGCTCTTCCGATCTNNVNVNVGTGATCTTTTTTTTTTTTTT TTTTTTTTTTV |
| 28S | GCCGGTAATACGACTCACTATAGGTCCCTACACGACGCTCTTCCGATCTNNVNVNVACAGTGTTTTTTTTTTTTTT TTTTTTTTTTV |
| 29S | GCCGGTAATACGACTCACTATAGGTCCCTACACGACGCTCTTCCGATCTNNVNVNVACCATGTTTTTTTTTTTTTT TTTTTTTTTTV |
| 30S | GCCGGTAATACGACTCACTATAGGTCCCTACACGACGCTCTTCCGATCTNNVNVNVACTCTGTTTTTTTTTTTTTT TTTTTTTTTTV |
| 31S | GCCGGTAATACGACTCACTATAGGTCCCTACACGACGCTCTTCCGATCTNNVNVNVACTCGATTTTTTTTTTTTTT TTTTTTTTTTV |
| 32S | GCCGGTAATACGACTCACTATAGGTCCCTACACGACGCTCTTCCGATCTNNVNVNVACGTACTTTTTTTTTTTTTT TTTTTTTTTTV |
| 33S | GCCGGTAATACGACTCACTATAGGTCCCTACACGACGCTCTTCCGATCTNNVNVNVACGTTGTTTTTTTTTTTTTT TTTTTTTTTTV |
| 34S | GCCGGTAATACGACTCACTATAGGTCCCTACACGACGCTCTTCCGATCTNNVNVNVACGTGATTTTTTTTTTTTTT TTTTTTTTTTV |

| | |
|---|---|
| 35S | GCCGGTAATACGACTCACTATAGGTCCCTACACGACGCTCTTCCGATCTNNVNVNVCTAGACTTTTTTTTTTTTTT TTTTTTTTTTV |
| 36S | GCCGGTAATACGACTCACTATAGGTCCCTACACGACGCTCTTCCGATCTNNVNVNVCTAGTGTTTTTTTTTTTTTT TTTTTTTTTTV |
| 37S | GCCGGTAATACGACTCACTATAGGTCCCTACACGACGCTCTTCCGATCTNNVNVNVCTAGGATTTTTTTTTTTTTT TTTTTTTTTTV |
| 38S | GCCGGTAATACGACTCACTATAGGTCCCTACACGACGCTCTTCCGATCTNNVNVNVCTCATGTTTTTTTTTTTTTT TTTTTTTTTTV |
| 39S | GCCGGTAATACGACTCACTATAGGTCCCTACACGACGCTCTTCCGATCTNNVNVNVCTCAGATTTTTTTTTTTTTT TTTTTTTTTTV |
| 40S | GCCGGTAATACGACTCACTATAGGTCCCTACACGACGCTCTTCCGATCTNNVNVNVCTTCGATTTTTTTTTTTTTT TTTTTTTTTTV |
| 41S | GCCGGTAATACGACTCACTATAGGTCCCTACACGACGCTCTTCCGATCTNNVNVNVCTGTACTTTTTTTTTTTTTT TTTTTTTTTTV |
| 42S | GCCGGTAATACGACTCACTATAGGTCCCTACACGACGCTCTTCCGATCTNNVNVNVCTGTGATTTTTTTTTTTTTT TTTTTTTTTTV |
| 43S | GCCGGTAATACGACTCACTATAGGTCCCTACACGACGCTCTTCCGATCTNNVNVNVTGAGACTTTTTTTTTTTTTT TTTTTTTTTTV |
| 44S | GCCGGTAATACGACTCACTATAGGTCCCTACACGACGCTCTTCCGATCTNNVNVNVTGCAACTTTTTTTTTTTTTT TTTTTTTTTTV |
| 45S | GCCGGTAATACGACTCACTATAGGTCCCTACACGACGCTCTTCCGATCTNNVNVNVTGCATGTTTTTTTTTTTTTT TTTTTTTTTTV |
| 46S | GCCGGTAATACGACTCACTATAGGTCCCTACACGACGCTCTTCCGATCTNNVNVNVTGCAGATTTTTTTTTTTTTT TTTTTTTTTTV |
| 47S | GCCGGTAATACGACTCACTATAGGTCCCTACACGACGCTCTTCCGATCTNNVNVNVTGTCACTTTTTTTTTTTTTT TTTTTTTTTTV |
| 48S | GCCGGTAATACGACTCACTATAGGTCCCTACACGACGCTCTTCCGATCTNNVNVNVTGTCGATTTTTTTTTTTTTT TTTTTTTTTTV |
| 49S | GCCGGTAATACGACTCACTATAGGTCCCTACACGACGCTCTTCCGATCTNNVNVNVTGGTACTTTTTTTTTTTTTT TTTTTTTTTTV |
| 50S | GCCGGTAATACGACTCACTATAGGTCCCTACACGACGCTCTTCCGATCTNNVNVNVGACATGTTTTTTTTTTTTTT TTTTTTTTTTV |
| 51S | GCCGGTAATACGACTCACTATAGGTCCCTACACGACGCTCTTCCGATCTNNVNVNVGATCACTTTTTTTTTTTTTT TTTTTTTTTTV |

| | |
|---|---|
| 52S | GCCGGTAATACGACTCACTATAGGTCCCTACACGACGCTCTTCCGATCTNNVNVNVGATCTGTTTTTTTTTTTTTTTTTTTTTTTTTV |
| 53S | GCCGGTAATACGACTCACTATAGGTCCCTACACGACGCTCTTCCGATCTNNVNVNVGATCGATTTTTTTTTTTTTTTTTTTTTTTTTV |
| 54S | GCCGGTAATACGACTCACTATAGGTCCCTACACGACGCTCTTCCGATCTNNVNVNVGAGTACTTTTTTTTTTTTTTTTTTTTTTTTTV |
| 55S | GCCGGTAATACGACTCACTATAGGTCCCTACACGACGCTCTTCCGATCTNNVNVNVAGACAGTTTTTTTTTTTTTTTTTTTTTTTTTV |
| 56S | GCCGGTAATACGACTCACTATAGGTCCCTACACGACGCTCTTCCGATCTNNVNVNVAGACCATTTTTTTTTTTTTTTTTTTTTTTTTV |
| 57S | GCCGGTAATACGACTCACTATAGGTCCCTACACGACGCTCTTCCGATCTNNVNVNVAGTGAGTTTTTTTTTTTTTTTTTTTTTTTTTV |
| 58S | GCCGGTAATACGACTCACTATAGGTCCCTACACGACGCTCTTCCGATCTNNVNVNVAGGAAGTTTTTTTTTTTTTTTTTTTTTTTTTV |
| 59S | GCCGGTAATACGACTCACTATAGGTCCCTACACGACGCTCTTCCGATCTNNVNVNVAGGACATTTTTTTTTTTTTTTTTTTTTTTTTV |
| 60S | GCCGGTAATACGACTCACTATAGGTCCCTACACGACGCTCTTCCGATCTNNVNVNVCAACAGTTTTTTTTTTTTTTTTTTTTTTTTTV |
| 61S | GCCGGTAATACGACTCACTATAGGTCCCTACACGACGCTCTTCCGATCTNNVNVNVCAACCATTTTTTTTTTTTTTTTTTTTTTTTTV |
| 62S | GCCGGTAATACGACTCACTATAGGTCCCTACACGACGCTCTTCCGATCTNNVNVNVCAACTCTTTTTTTTTTTTTTTTTTTTTTTTTV |
| 63S | GCCGGTAATACGACTCACTATAGGTCCCTACACGACGCTCTTCCGATCTNNVNVNVCACTCATTTTTTTTTTTTTTTTTTTTTTTTTV |
| 64S | GCCGGTAATACGACTCACTATAGGTCCCTACACGACGCTCTTCCGATCTNNVNVNVCACTTCTTTTTTTTTTTTTTTTTTTTTTTTTV |
| 65S | GCCGGTAATACGACTCACTATAGGTCCCTACACGACGCTCTTCCGATCTNNVNVNVCAGAAGTTTTTTTTTTTTTTTTTTTTTTTTTV |
| 66S | GCCGGTAATACGACTCACTATAGGTCCCTACACGACGCTCTTCCGATCTNNVNVNVCAGACATTTTTTTTTTTTTTTTTTTTTTTTTV |
| 67S | GCCGGTAATACGACTCACTATAGGTCCCTACACGACGCTCTTCCGATCTNNVNVNVTCACCATTTTTTTTTTTTTTTTTTTTTTTTTV |
| 68S | GCCGGTAATACGACTCACTATAGGTCCCTACACGACGCTCTTCCGATCTNNVNVNVTCACTCTTTTTTTTTTTTTTTTTTTTTTTTTV |

| | |
|---|---|
| 69S | GCCGGTAATACGACTCACTATAGGTCCCTACACGACGCTCTTCCGATCTNNVNVNVTCCTCATTTTTTTTTTTTTT TTTTTTTTTTV |
| 70S | GCCGGTAATACGACTCACTATAGGTCCCTACACGACGCTCTTCCGATCTNNVNVNVTCCTTCTTTTTTTTTTTTTT TTTTTTTTTTV |
| 71S | GCCGGTAATACGACTCACTATAGGTCCCTACACGACGCTCTTCCGATCTNNVNVNVTCTGTCTTTTTTTTTTTTTT TTTTTTTTTTV |
| 72S | GCCGGTAATACGACTCACTATAGGTCCCTACACGACGCTCTTCCGATCTNNVNVNVGTCTTCTTTTTTTTTTTTTT TTTTTTTTTTV |
| 73S | GCCGGTAATACGACTCACTATAGGTCCCTACACGACGCTCTTCCGATCTNNVNVNVGTTGAGTTTTTTTTTTTTTT TTTTTTTTTTV |
| 74S | GCCGGTAATACGACTCACTATAGGTCCCTACACGACGCTCTTCCGATCTNNVNVNVGTTGTCTTTTTTTTTTTTTT TTTTTTTTTTV |
| 75S | GCCGGTAATACGACTCACTATAGGTCCCTACACGACGCTCTTCCGATCTNNVNVNVGTGAAGTTTTTTTTTTTTTT TTTTTTTTTTV |
| 76S | GCCGGTAATACGACTCACTATAGGTCCCTACACGACGCTCTTCCGATCTNNVNVNVACAGACTTTTTTTTTTTTTT TTTTTTTTTTV |
| 77S | GCCGGTAATACGACTCACTATAGGTCCCTACACGACGCTCTTCCGATCTNNVNVNVACAGGATTTTTTTTTTTTTT TTTTTTTTTTV |
| 78S | GCCGGTAATACGACTCACTATAGGTCCCTACACGACGCTCTTCCGATCTNNVNVNVACCAACTTTTTTTTTTTTTT TTTTTTTTTTV |
| 79S | GCCGGTAATACGACTCACTATAGGTCCCTACACGACGCTCTTCCGATCTNNVNVNVACCAGATTTTTTTTTTTTTT TTTTTTTTTTV |
| 80S | GCCGGTAATACGACTCACTATAGGTCCCTACACGACGCTCTTCCGATCTNNVNVNVACTCACTTTTTTTTTTTTTT TTTTTTTTTTV |
| 81S | GCCGGTAATACGACTCACTATAGGTCCCTACACGACGCTCTTCCGATCTNNVNVNVCTCAACTTTTTTTTTTTTTT TTTTTTTTTTV |
| 82S | GCCGGTAATACGACTCACTATAGGTCCCTACACGACGCTCTTCCGATCTNNVNVNVCTTCACTTTTTTTTTTTTTT TTTTTTTTTTV |
| 83S | GCCGGTAATACGACTCACTATAGGTCCCTACACGACGCTCTTCCGATCTNNVNVNVCTTCTGTTTTTTTTTTTTTT TTTTTTTTTTV |
| 84S | GCCGGTAATACGACTCACTATAGGTCCCTACACGACGCTCTTCCGATCTNNVNVNVCTGTTGTTTTTTTTTTTTTT TTTTTTTTTTV |
| 85S | GCCGGTAATACGACTCACTATAGGTCCCTACACGACGCTCTTCCGATCTNNVNVNVTGAGTGTTTTTTTTTTTTTT TTTTTTTTTTV |

| | |
|---|---|
| 86S | GCCGGTAATACGACTCACTATAGGTCCCTACACGACGCTCTTCCGATCTNNVNVNVTGAGGATTTTTTTTTTTTTT TTTTTTTTTTV |
| 87S | GCCGGTAATACGACTCACTATAGGTCCCTACACGACGCTCTTCCGATCTNNVNVNVTGTCTGTTTTTTTTTTTTTT TTTTTTTTTTV |
| 88S | GCCGGTAATACGACTCACTATAGGTCCCTACACGACGCTCTTCCGATCTNNVNVNVTGGTTGTTTTTTTTTTTTTT TTTTTTTTTTV |
| 89S | GCCGGTAATACGACTCACTATAGGTCCCTACACGACGCTCTTCCGATCTNNVNVNVTGGTGATTTTTTTTTTTTTT TTTTTTTTTTV |
| 90S | GCCGGTAATACGACTCACTATAGGTCCCTACACGACGCTCTTCCGATCTNNVNVNVGAAGACTTTTTTTTTTTTTT TTTTTTTTTTV |
| 91S | GCCGGTAATACGACTCACTATAGGTCCCTACACGACGCTCTTCCGATCTNNVNVNVGAAGTGTTTTTTTTTTTTTT TTTTTTTTTTV |
| 92S | GCCGGTAATACGACTCACTATAGGTCCCTACACGACGCTCTTCCGATCTNNVNVNVGAAGGATTTTTTTTTTTTTT TTTTTTTTTTV |
| 93S | GCCGGTAATACGACTCACTATAGGTCCCTACACGACGCTCTTCCGATCTNNVNVNVGACAACTTTTTTTTTTTTTT TTTTTTTTTTV |
| 94S | GCCGGTAATACGACTCACTATAGGTCCCTACACGACGCTCTTCCGATCTNNVNVNVGACAGATTTTTTTTTTTTTT TTTTTTTTTTV |
| 95S | GCCGGTAATACGACTCACTATAGGTCCCTACACGACGCTCTTCCGATCTNNVNVNVGAGTTGTTTTTTTTTTTTTT TTTTTTTTTTV |
| 96S | GCCGGTAATACGACTCACTATAGGTCCCTACACGACGCTCTTCCGATCTNNVNVNVGAGTGATTTTTTTTTTTTTT TTTTTTTTTTV |

Other primers

The RandomhexRT primer (**Table 2**) serves as the primer for the RT2 step (see below) in our protocol. For this step, we have modified the adaptor sequence (highlighted in yellow) to match the 3' end of the Illumina Index PCR Primers.

The P5 primer starts with Illumina P5 sequences (colored in red) and is followed by adaptor sequences that overlap with the adaptor sequences in the barcoding primer. In turn, the index primer begins with the Illumina P7 sequence and is followed by a 6-nt index sequence (highlighted in green) and adaptor sequences (highlighted in yellow) that overlap with the adaptor sequences in the RandomhexRT primer.

The RandomhexRT primer was purified by high-performance liquid chromatography (HPLC), while the other primers were purified using PAGE. All the primers were synthesized by Sangon Biotech.

**Table 2. Other primers**

| | |
|---|---|
| RandomhexRT | GTTCAGACGTGTGCTCTTCCGATCTNNNNNN |
| P5 | AATGATACGGCGACCACCGAGATCTACACACTCTTTCCCTACACGACGCTCTTCC |
| Index-1 | CAAGCAGAAGACGGCATACGAGATCGTGATGTGACTGGAGTTCAGACGTGTGCTCTTCC |
| Index-2 | CAAGCAGAAGACGGCATACGAGATACATCGGTGACTGGAGTTCAGACGTGTGCTCTTCC |
| Index-5 | CAAGCAGAAGACGGCATACGAGATCACTGTGTGACTGGAGTTCAGACGTGTGCTCTTCC |
| Index-7 | CAAGCAGAAGACGGCATACGAGATGATCTGGTGACTGGAGTTCAGACGTGTGCTCTTCC |
| Index-4 | CAAGCAGAAGACGGCATACGAGATTGGTCAGTGACTGGAGTTCAGACGTGTGCTCTTCC |
| Index-6 | CAAGCAGAAGACGGCATACGAGATATTGGCGTGACTGGAGTTCAGACGTGTGCTCTTCC |
| Index-8 | CAAGCAGAAGACGGCATACGAGATTCAAGTGTGACTGGAGTTCAGACGTGTGCTCTTCC |

Method

1. Prior to constructing the sequencing library, dilute all RNA samples to 25 ng/μL. Next, take 5 μL of several random samples to run on a 1% agarose gel in TBE buffer to check the RNA quality.

   Here is an example:

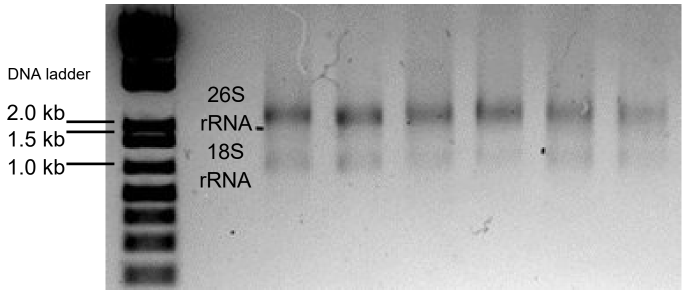


   For a total RNA sample with good quality, you will see two main bands, the upper 26S rRNA band and the lower 18S rRNA band, without obvious signs of degradation. If RNA is degraded, you will see bands or smears under 1.0 kb on the DNA ladder.

2. Begin library construction by performing **first strand reverse transcription** (RT1). This step barcodes the samples and needs to be done in 96-well PCR plate. Perform the experiment following the steps below.
    a. In a 96-well PCR plate, add the following reagents to each well:
        - Total RNA (25 ng/μL): 2 μL
        - Barcoding primer (2.5 μM): 2 μL
        - dNTP mix (dilute to 2.5 mM of each): 2 μL
    b. Spin down the 96-well plate and incubate the mixture at 65℃ for 5 min, and then cool down on ice for 2 min.
    c. Mix the following reagents and add 5 μL of the mix to each well of the 96-well plate:
        - First Strand Buffer: 220 μL
        - DTT (0.1 M): 50 μL
        - RNaseOUT: 30 μL
        - SuperScript™ III: 10 μL
        - $H_2O$: 200 μL
    d. Spin down the 96-well plate and incubate the plate at 50℃ for 90 min; then 70℃ for another 20 min.

    ***Checkpoint**: keep the products at 4℃ for several hours, or -80℃ for several months.*

3. Pool half of the RT1 products for each sample into Eppendorf tubes while keeping everything on ice.

    We recommend pooling ~50 samples to one sequencing library (tube). After pooling, purify the pooled library with 1.2X AMPure XP beads following the original CEL-Seq2 protocol [5], hereinafter follow the same protocol for beads purification). Elute the beads with 25 μL $H_2O$ for each library. This produces 25 μL RT1 product for each pooled library.

4. **Exonuclease I digestion**: this step is to remove residual primers from the first reverse transcription step.

   For each library, mix the following reagents on ice:

   - RT1 product: 25 μL
   - 10X digestion buffer: 3 μL
   - Exonuclease I: 2 μL

   Then briefly vortex the mixture and incubate it at 37℃ for 30 min. Purify the products with 1.2X AMPure XP beads. Elute the beads with 25 μL $H_2O$ for each library.

5. Perform the **second strand cDNA synthesis** with the following steps.
   a. For each library, mix the following reagents on ice:
      - Elution from step 4: 22.2 μL
      - 5X second strand reaction buffer: 6 μL
      - dNTP mix (10 mM): 0.6 μL
      - *E. coli* DNA ligase (10 u/μL): 0.2 μL
      - DNA polymerase I (10 u/μL): 0.8 μL
      - Ribonuclease H (2 u/μL): 0.2 μL
   b. Then briefly vortex the mixture and incubate it at 16℃ for 2 hours with lid open and then at 65℃ for 20 min with lid heated at > 80℃.

      ***Checkpoint****: keep the products at 4℃ for several hours, or -80℃ for several months.*
   c. Purify the products with 1.2X AMPure XP beads. Elute the beads with 5 μL $H_2O$ for each library.

   **Quality Control**: Take 1 μL purified product to measure the concentration using the Qubit dsDNA HS (High Sensitivity) Assay Kit. Here is an example of the results of 6 libraries in one batch:

| | Lib1 | Lib 2 | Lib 3 | Lib 4 | Lib 5 | Lib 6 |
|---|---|---|---|---|---|---|
| Concentration (ng/μL) | 2.16 | 2.14 | 2.71 | 2.52 | 2.38 | 2.60 |

*We are not sure if the concentration reflects the real concentration of dsDNA, but the measured concentration positively correlated with the RNA yield in the following In vitro transcription step. We recommend a minimum concentration of 1 ng/μL for a library pooled with ~50 samples to proceed to the next step.*

6. ***In vitro* transcription** (IVT). Use HiScribe™ T7 High Yield RNA Synthesis Kit to perform IVT with the following steps.

   a. For each library mix the following reagents at room temperature:
      - Product from step 5: 4 μL
      - NTPs: 1 μL of each (4 μL total)
      - 10X IVT buffer: 1 μL
      - T7 enzyme: 1 μL

      Briefly vortex the mixture and incubate it at 37℃ overnight in a PCR machine.

   b. Purify RNA using 1.8X RNAClean XP beads. Elute the beads with 30 μL $H_2O$ for each library.

**Quality Control:** measure the product concentration by NanoDrop. Here is an example of the results of 6 libraries in one batch:

| | Lib1 | Lib 2 | Lib 3 | Lib 4 | Lib 5 | Lib 6 |
|---|---|---|---|---|---|---|
| Concentration (ng/μL) | 572 | 660 | 627 | 529 | 532 | 582 |

*For most libraries in our hands, the concentration of IVT products were more than 300 ng/μL. We recommend a minimal concentration of 100 ng/μL for a library pooled with ~50 samples to proceed to the next step.*

c. Diute all IVT products to 300 ng/μL.

7. **Fragmentation**: this step is to fragment the IVT RNA products.
   a. For each library mix the following reagents on ice:
      - IVT RNA products: 4 μL (1.2 μg in total. If IVT RNA products concentration is less than 300 ng/μL, increase the volume to make the total input to be ~1.2 μg)
      - $H_2O$: 5 μL (If IVT RNA products concentration is less than 300 ng/μL, decrease the volume to make the total volume of IVT RNA products and $H_2O$ be 9 μL)
      - 10X Fragmentation Buffer: 1 μL
   b. Briefly vortex the mixture and incubate at 70℃ for 2 min.
   c. Add 1 μL of the Stop Solution and immediately proceed to purify the products with ice cold 1.8X RNA Clean XP beads. Elute the beads with 20 μL $H_2O$ for each library.

**Quality Control 1:** measure the product concentration by NanoDrop. Here is an example of the results of 6 libraries in one batch:

| | Lib1 | Lib 2 | Lib 3 | Lib 4 | Lib 5 | Lib 6 |
|---|---|---|---|---|---|---|
| Concentration (ng/μL) | 47.8 | 41.8 | 46.8 | 46.2 | 44.4 | 39.0 |

*The concentration of fragmented products was usually more than 30 ng/µL in our hands. The recovery rate is >50% and ~70% for most cases.*

d. Dilute all samples to 25 ng/µL.

**Quality Control 2:** Run a 1.2% TBE gel using 1.5 µL diluted IVT products or 5 µL diluted fragmentation product to check RNA size. Here is a gel image of 6 IVT products (left to right) before (larger-size band) and after (smaller-size band) fragmentation.

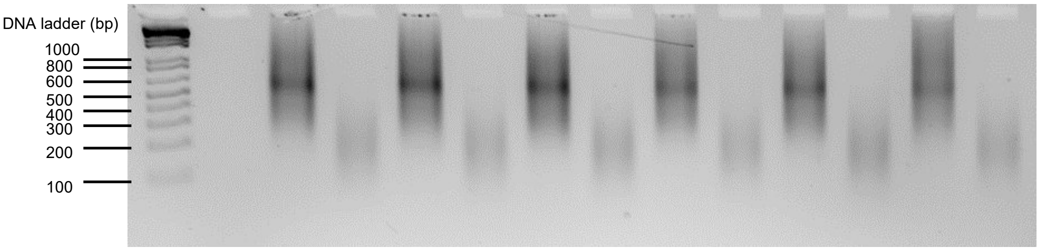


*The size of IVT and Fragmentation products were very stable in our libraries. The main band of IVT product is at the position between 500 bp to 600 bp of the DNA ladder while the Fragmentation product is at the position between 100 bp to 200 bp of DNA ladder. A smaller IVT product size indicates an inefficient RT1 reaction. A larger Fragmentation product size indicates inefficient fragmentation, while smaller Fragmentation product size indicates over fragmentation. (Please be aware that the DNA ladder only provides a reference for the relative fragment size; it does not indicate the actual sizes of RNA fragments.)*

8. **The second reverse transcription** (RT2): this step is to generate cDNA from the fragmented RNA and add adaptor sequence to the cDNA product.
    a. For each library mix the following reagents on ice:
        - Fragmented RNA (25 ng/µL): 3 µL
        - RandomhexRT primer (10 µM): 2 µL

- dNTP mix (10 mM of each): 1 μL
- $H_2O$: 7 μL

b. briefly vortex the mixture and incubate it at 65℃ for 5 min and then cool on ice for 2 min.

c. add the following reagents to each library reaction on ice:
   - First Strand Buffer: 4 μL
   - DTT (0.1 M): 2 μL
   - RNaseOUT: 0.5 μL
   - SuperScript™ III: 0.5 μL

d. briefly vortex the mixture and incubate it with the following temperature in a PCR machine:
   - 25 ℃ 10 min,
   - 50 ℃ 60 min,
   - 70 ℃ 15min.

e. Purify the products with 1.2X AMPure XP beads. Elute the beads with 20 μL $H_2O$ for each library.

9. **PCR amplification**: this is the final step for WPS library construction and produces the final library for sequencing.
   a. For each library mix the following reagents on ice:
      - RT2 product: 10 μL
      - $H_2O$: 8 μL
      - Index primer primer (10 μM): 1 μL
      - P5 primer (10 μM): 1 μL
      - Kapa HIFI 2X mix: 20 μL

   b. briefly vortex the mixture, then do PCR using the following cycling conditions:
      - 98 ℃ 2 min,
      - 12 cycles of
        1) [98 ℃ 10 s
        2) 60 ℃ 30 s

3) 72 ℃ 30 s],

- 72 ℃ 5 min,
- Hold at 4 ℃.

c. Use 1X AMPure XP beads to purify the PCR products. Elute the beads with 25 μL $H_2O$ for each library.
d. Use 1X AMPure XP beads to purify the 25 μL eluent from step c. Elute the beads with 25 μL $H_2O$ for each library.

**Quality Control:** measure the library concentration by Qubit dsDNA HS (High Sensitivity) Assay Kit. Here is an example of the results of 6 libraries in one batch:

| | Lib1 | Lib 2 | Lib 3 | Lib 4 | Lib 5 | Lib 6 |
|---|---|---|---|---|---|---|
| Concentration (ng/μL) | 11.6 | 12.7 | 13.4 | 12.3 | 12.6 | 12.4 |

*The concentration of libraries is usually ~10 ng/μL. >5 ng/μL is acceptable.*

10. Pre-sequencing quality control of final libraries

*This step is to check the size distribution of the final library. We use Qubit to quantify concentrations of libraries and use one of the following two methods to check the size distribution of each library.*

Method 1: Fragment analyzer

Mix 1 μL library and 2 μL $H_2O$ and run Fragment analyzer with the 3 μL solution. Here is an example of fragment analyzer result of a good library:

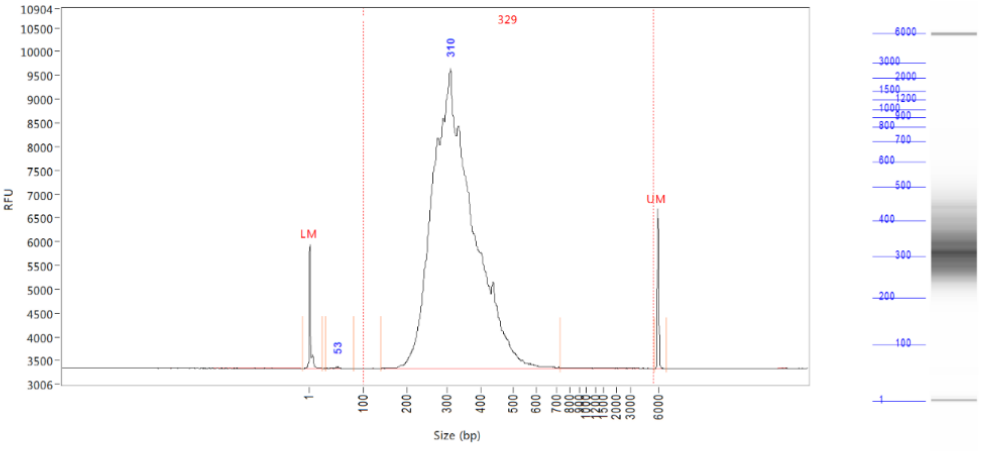


| Peak | Size (bp) | Conc. (ng/uL) | From (bp) | To (bp) | Avg. Size (bp) | CV% | RFU | Corr. Peak Area |
|---|---|---|---|---|---|---|---|---|
| 1 | 1 (LM) | 0.0152 | 0 | 26 | 2 | 172.70 | 2609 | 16.540 |
| 2 | 53 | 0.0054 | 31 | 84 | 50 | 8.80 | 36 | 0.490 |
| 3 | 310 | 8.2779 | 140 | 728 | 327 | 19.99 | 6304 | 748.240 |
| 4 | 6000 (UM) | 0.0091 | 5716 | 6873 | 5995 | 1.30 | 3356 | 9.871 |

| | | |
|---|---|---|
| TIC: | 8.2834 | ng/uL |
| TIM: | 41.822 | nmole/L |
| Total Conc.: | 8.2992 | ng/uL |

Smear Analysis 100 bp to 5500 bp 8.2937 ng/ul 99.9 %Total 41.408 nmole/L 329 Avg. Size (b.p.) 27.57 %CV

*The peak size is expected to be ~300 bp with peak expansion from 200 bp to ~500 bp. Any obvious shift from this range indicates problems in previous steps (e.g., RT1 and fragmentation) and may cause trouble in the sequencing step.*

Method 2: Gel electrophoresis

We found that it is robust enough to simply run ~30 ng or more of the final library on a 1.2% TBE gel for quality control of the fragment size. This is more cost efficient and faster than the Fragment analyzer that requires a special facility. Here is an example of the results of 6 libraries from one batch:

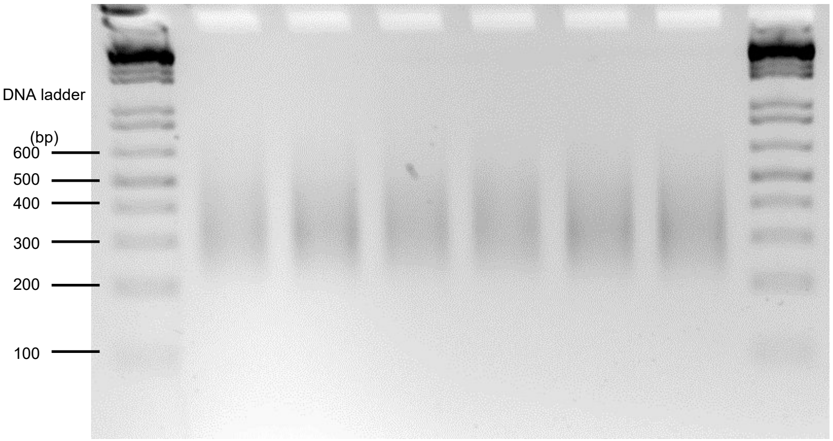

*We expect the main band spread to be between 200-500 bp with a peak at ~300 bp. Any obvious shift from this range needs to be further diagnosed by fragment analyzer or resolved by remaking the library.*

**Section IV: Data analysis**

Please refer to [https://github.com/XuhangLi/WPS](https://github.com/XuhangLi/WPS) for a full tutorial of WPS data analysis.

Notes on the use of same-day vector controls

As discussed in the previous sections, standard WPS protocol prepares two independent cultures of vector control in each experiment, which we refer to as same-day replicates. This is in principle different than the stringent biological replicates that are done in different days. Interestingly, we found that the level of gene expression correlation between same-day replicates closely aligned with that between different-day replicates (see the figure below). Therefore, we used both same-day and different-day replicates in our DE analysis

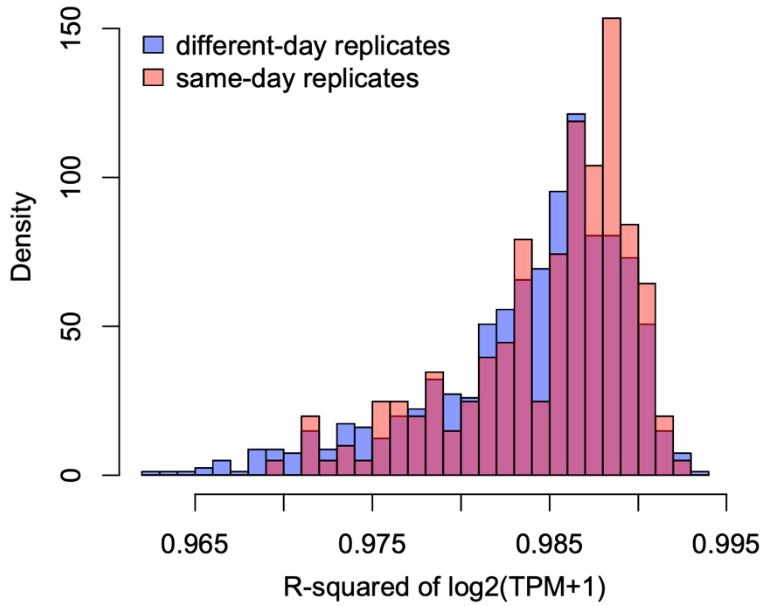


to increase power. We recommend that users inspect their own data to determine if both replicates should be included. Specifically, please (1) confirm that a similar closely aligned variance level is observed, and (2) test DE analysis using either both types of replicates or only the different-day replicates and check the differences in the results. Based on our experience, DE analysis using both replicates in fact gives more conservative results (less DEGs) in many conditions.
